# Resilience to neuronal hyperactivity and restoration of the neuroimmune interactome and decision-making by blocking fibrin in a model of Alzheimer’s disease

**DOI:** 10.64898/2026.05.01.722077

**Authors:** Zhaoqi Yan, Andrew S. Mendiola, Kelli Lauderdale, Keun-Young Kim, Yu Yong, Kun Leng, Eric A. Bushong, Renaud Schuck, Anke Meyer-Franke, Ayushi Agrawal, Michela Traglia, Natalie Gill, Reuben Thomas, Jonah N. Keller, Nikolaos Karvelas, Miguel F Vasquez, Daniel C Ballard, Matthew Madany, Jeffrey Simms, Brandon Guo, Reshmi Tognatta, Maria del Pilar S. Alzamora, Rosa Meza-Acevedo, Belinda Cabriga, Kellie N. Pierson, Lida Kourita, Kayoung Han, Jae K. Ryu, Bruce L. Miller, Fanny M. Elahi, Mark H. Ellisman, Jorge J. Palop, Katerina Akassoglou

**Author notes:** These authors contributed equally (Zhaoqi Yan, Andrew S. Mendiola). Correspondence (K.A.).

## Abstract

Cerebrovascular pathology and neuronal network dysfunction are early features of Alzheimer’s disease (AD) associated with neuroinflammation and cognitive decline, but the vascular and immune triggers of neuronal hyperactivity remain largely unknown. Here, we show that the blood coagulation protein fibrin disrupts microglia-neuron interactions, promoting neuronal hyperactivity in an AD mouse model. Genetic elimination of the fibrin inflammatory domain reduced neuronal hyperactivity, restored dynamic microglial interactions with active neurons and protected from high-risk decision making in 5XFAD mice. Leveraging the transcriptional signatures of microglia and inhibitory and excitatory neurons, a ligand–receptor atlas revealed fibrin-dependent disruption of innate immune and glutamatergic signaling between microglia and neurons in AD mice. Patients with AD also showed a correlation of cerebrospinal fluid (CSF) fibrinogen levels with biomarkers of inflammation, vascular and synaptic dysfunction. Thus, resilience to neuronal hyperactivity and restoration of the neuroimmune interactome by targeting fibrin may have therapeutic implications for Alzheimer’s disease and related conditions. There is a companion manuscript submitted to bioRxiv (Lauderdale et al., 2026)

**Highlights:**

- Vascular-microglia axis drives neuronal hyperactivity
- Fibrin inflammatory activity disrupts the microglia-neuron interactome
- Microglia activation by fibrin impairs decision-making in AD mice
- Synaptic dysfunction and immune biomarkers correlate with CSF fibrinogen in AD patients

## Introduction

Neuronal network dysfunction and cerebrovascular pathology are early features of Alzheimer’s disease (AD).^1–5^ Cerebrovascular dysfunction is a major risk factor for dementia and synaptic dysfunction in aging and AD.^6–8^ Compromised vascular integrity and blood-brain barrier (BBB) disruption results in the influx of blood proteins into the CNS detected in mild cognitive impairment (MCI) as well as early AD and correlates with disease progression.^2,9–13^ Neural oscillations underlie learning, memory, and cognitive function, and aberrant neuronal activity is increased early in AD and in related animal models.^14–16^ AD patients with subclinical epileptiform activity exhibit network hyperexcitability, which is associated with accelerated cognitive impairment in human AD and mouse models.^3,4,17–20^ Excessive neuronal network activation can also manifest as epilepsy and BBB dysfunction positively correlates with epileptogenesis and seizure recurrence in patients with epilepsy.^21–23^ Microglia regulate neuronal activity in both physiology and epilepsy models^24–27^. However, the role of microglia in neuronal hyperactivity in AD and the molecular triggers that disrupt microglia-neuron interactions remain poorly understood. Soluble Aβ, tau protein, and structural axonal defects regulate neuronal hyperactivity in AD, ^16,28–30^ but the role of vascular and neuroimmune mechanisms in the AD brain driving neuronal network abnormalities are largely unknown.

BBB disruption and intracerebral microhemorrhages/microbleeds are manifestations of vascular damage in individuals with AD and correlate with the progression from mild cognitive impairment to AD.^2,31^ Following BBB disruption or vascular injury, the blood coagulation factor fibrinogen is deposited as insoluble fibrin in the AD brain.^32–36^ Fibrin deposition correlates with disease progression and neurological dysfunction in patients with AD, as shown by epidemiological, neuropathological, and biomarker studies.^2,9–11,13,37–42^ Fibrin co-localizes with activated microglia, Aβ plaques, and axonal spheroids in AD brains.^33,34,43,44^ Conversion of fibrinogen to fibrin exposes an inflammatory fibrin domain that selectively induces neurotoxic microglia responses via CD11b/CD18 receptor binding and mechanotransduction signaling, including increased oxidative stress, pro-inflammatory cytokine release, and synapse elimination.^12,34,43,45,46^ Genetic inactivation of the fibrin inflammatory domain protects AD mice from neuronal loss, neurotoxic microglia reactivity, cognitive impairment in learning and memory tests and spontaneous behavioral alterations.^34,45,47^ However, despite this evidence for a dynamic interplay between cerebrovascular dysfunction and neuroinflammation, ^12,46,48^ the role of fibrin in neuronal hyperactivity remains largely unknown.

Here we provide evidence for a fundamental role for fibrin in driving neuronal hyperactivity and impaired decision-making by disrupting microglia functions within neuronal circuits, highlighting the therapeutic potential of targeting fibrin in AD.

## Results

### Fibrin drives microglia-mediated neuronal hyperactivity

Neuronal hyperactivity is prominent in models of AD,^16,28,30^ but the role of neurovascular dysfunction has not been defined. To examine the effect of fibrin inflammatory signaling on neuronal activity, we crossed 5XFAD mice with *Fgg^γ390-396A^* mice, which express a mutant fibrinogen γ chain lacking the fibrin-CD11b binding domain while retaining normal clotting function.^34,45^ We measured neuron-specific spontaneous Ca^2+^ transients in awake, head-fixed 5XFAD mice injected with AAV-jRCaMP1b by *in vivo* two-photon (2P) time-lapse imaging (Figure 1A). 5XFAD mice exhibited an increase of hyperactive neurons, defined as neurons with increased frequency of Ca^2+^ transients (Figure 1B) consistent with prior findings.^28^ Genetic inhibition of the fibrin inflammatory domain normalized neuronal hyperactivity in 5XFAD mice to non-transgenic (NTG) level (Figure 1B). Quantification of synchronized neuronal firing revealed in 5XFAD mice increased neuronal hypersynchrony, which was rescued in 5XFAD:*Fgg^γ390-396A^* mice to NTG levels (Figure 1C), suggesting that fibrin-microglia interactions promote synchronized network activity.

**Figure 1.**
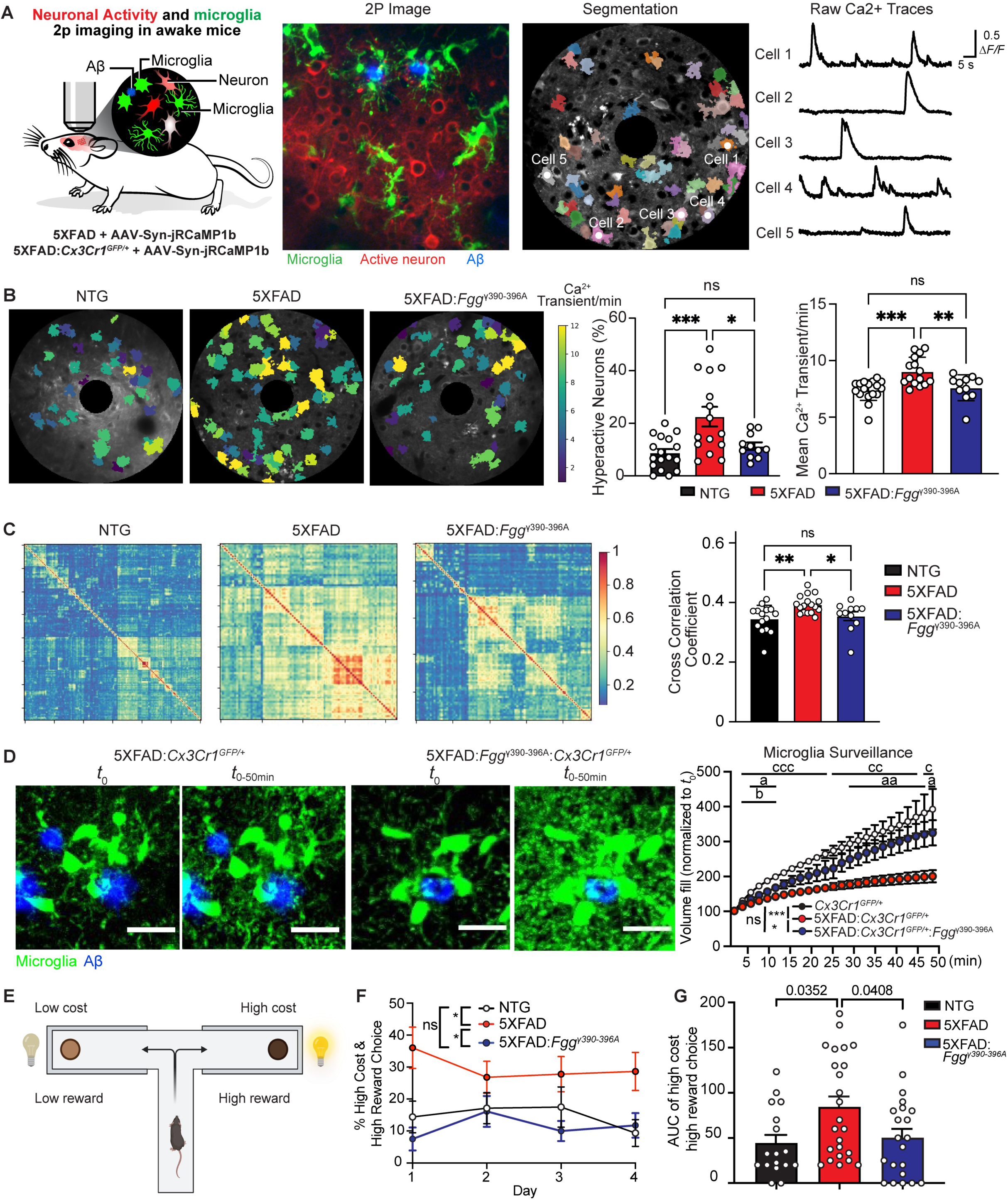
Fibrin induces microglia-mediated neuronal hyperactivity and reduces high-risk behavior in AD mice. (A) Left, Schematic of in vivo 2P imaging for neuronal activity and microglia dynamic analysis in awake mice. Middle, representative 2P image of simultaneous recording of neuronal activity (jRCaMP1b, red), microglia (GFP, green) and amyloid plaque (Methoxy-X04, blue) from cortex. Right, example of neuron segmentation by CaImAn and raw Ca^2+^ traces from individual cortical neurons. (B) Representative heatmaps of cumulative spontaneous awake neuronal Ca^2+^ transient and quantification of % of hyperactive neurons and Mean Ca^2+^ transient/min for all cells. Each dot represents one animal with combined neurons from 2-3 ROI. Data are shown as mean ± s.e.m. n = 17 NTG, n = 15 5XFAD, and n = 11 5XFAD: *Fgg^γ390-396A^*. *P < 0.05 or ***P < 0.001 by one-way ANOVA with Tukey’s multiple comparisons test. ns, no significance. (C) Clustered cross-correlation matrices of Ca^2+^ transients (left) to assess neuronal synchrony and quantification of the cross-correlation coefficient (right). Each dot represents one animal. Data are shown as mean ± s.e.m. n = 17 NTG, n = 15 5XFAD, and n = 11 5XFAD: *Fgg^γ390-396A^*. *P < 0.05 or **P < 0.01 by one-way ANOVA with Tukey’s multiple comparisons test. ns, no significance. (D) *In vivo* 2P time-lapse imaging (left) and quantification (right) of cumulative microglial (green) surveillance near amyloid plaque (blue) over time. Scale bar, 20 µm. Data are shown as the mean ± s.e.m. n = 6 NTG, *n* = 9 5XFAD, n = 5 5XFAD: *Fgg^γ390-396A^* mice. Genotype effect over time *\*\*\*P* < 0.001, * *P* < 0.05 by unpaired two-sample t-test of the mean AUC. a (5XFAD: *Fgg^γ390-396A^* vs 5XFAD), b (NTG vs 5XFAD: *Fgg^γ390-396A^*), c (NTG vs 5XFAD) indicates *P* at individual time points by permutation test. a, b, c *P* < 0.05 or aa,cc *P* < 0.01, ccc, *P* < 0.001. (E-G) Assessment of risk-taking behavior by the Cost-Benefit Conflict (CBC) Test. (E) Schematic of the CBC Test. (F) Quantification of the percentages of high-cost and high-reward choices (pure chocolate milk with bright light on the left) over 4 days (mean ± s,e.m.) and (G) area under the curve of high-cost high-reward choices (mean ± s.e.m.). Data are from n = 17 NTG, n = 24 5XFAD, and n = 22 5XFAD:*Fgg^γ390-396A^* mice. \**P* < 0.05 by (F) repeated-measure one-way ANOVA and (G) one-way ANOVA with Tukey’s multiple comparisons test.

Since impaired microglial dynamics can promote neuronal hyperexcitability,^27^ we next used *in vivo* 2P imaging to determine the role of fibrin inflammatory signaling in microglial surveillance in AD mice. Surveillance of microglia surrounding Aβ plaques was significantly reduced in 5XFAD:*Cx3cr1^GFP/+^* mice compared to age-matched *Cx3cr1^GFP/+^* controls, but was restored in 5XFAD:*Fgg^γ390-396A^*:*Cx3cr1^GFP/+^* mice (Figures 1D and S1A, Supplementary Video 1). In addition, process tip, velocity and rates of process protraction and retraction of microglial surrounding Aβ plaques were significantly improved in 5XFAD:*Fgg^γ390-396A^*:C*x3cr1^GFP/+^* mice compared with 5XFAD:*Cx3cr1^GFP/+^* mice (Figure S1B-E). Using 3D-Morph morphology analysis, plaque-associated microglia in 5XFAD:*Fgg^γ390-396A^*mice were more ramified, with a larger cell volume and with more branch points than microglia in 5XFAD:*Cx3cr1^GFP/+^* mice (Figure S1F-I), suggesting that inhibition of the fibrin inflammatory domain restores microglial brain surveillance in amyloid pathology. Together, these results show that fibrin inflammatory signaling is required for impaired microglial surveillance and neuronal hyperactivity in AD mice.

### Fibrin increases risky decsion-making in AD mice

Increased risk-taking and impaired decision-making are recognized as an early indicator of executive dysfunction in AD that may precede memery loss, also observed in patients with epilepsy.^49–54^ We hypothesized that fibrin-induced microglial-mediated neuronal hyperactivity may dysregulate neuronal circuits involved in decision-making. We utilized a Cost-Benefit Conflict (CBC) task assessing non-optimal high risk decision-making by measuring how rodents resolve motivational conflict by weighing a high-value reward (pure chocolate milk) paired with a high cost (bright, aversive light) against a low reward (diluted chocolate milk) paired with a low cost (dim light). ^55,56^ We first habituated and trained in forced-entry trials to associate a high reward (100% chocolate milk) with bright light and a low reward (80% chocolate milk) with dim light, followed by free-choice trials to record their decisions (Figure 1E). Compared to NTG controls, 5XFAD mice exhibited increased high cost/high benefit choices (Figure 1F-G), consistent with previously reported high risk decision in foraging/predator behavioral tests^57^. In contrast, 5XFAD:*Fgg^γ390-396A^* mice had reduced high cost/high benefit choices than 5XFAD mice similar to NTG controls (Figure 1F-G), suggesting that fibrin drives non-optimal decision-making. No differences were observed in the time spent in dark and light without a reward or in the average number of entries per trial in the presence of a reward, suggesting that the higher percent of risk choices were not due to preferences in light exposure or locomotor activity (Figure S1J, K). Together, these data suggest that inhibition of fibrin-microglia activation may improve optimal decision-making in AD.

### Genetic blockade of the fibrin inflammatory domain restores microglia contacts with neurons in AD mice

Direct junctional contacts between microglial processes and neuronal somata suppress aberrant neuronal activity in the healthy brain.^27,58^ We employed a correlative multimodal multiscale microscopy pipeline to study the ultrastructural features of dynamic microglia-neuron interactions around Aβ plaques (Figure S2). By *in vivo* 2P imaging of microglia (GFP) and active neurons (jRCaMP1b) near Aβ plaques (methoxy-X04), we found increased microglia contacts with neuronal somata with closer proximity of microglia processes and neurons in 5XFAD:*Fgg^γ390–396A^:Cx3cr1^GFP/+^* compared to 5XFAD:*Cx3cr1^GFP/+^* mice (Figure 2A). Quantification of dynamic microglia motility in awake animals revealed increased microglial processes around active neurons in 5XFAD:*Fgg^γ390–396A^:Cx3cr1^GFP/+^* mice than in 5XFAD mice near Aβ (Figure 2B). To understand the subcellular dynamics underlying microglia-mediated regulation of the neuronal connectome in AD, we performed correlative light and electron microscopy (CLEM) by co-registering confocal microscopy imaging volumes of microglia, neurons and Aβ plaques with serial block-face electron microscopy (SBEM) using X-ray microscopy (XRM).^27^ This approach enabled precise ultrastructural mapping of microglial processes in relation to neuronal compartments within the Aβ plaque microenvironment (Figure S2). For semi-automatic segmentation to facilitate large-scale mapping of microglia-neuron interactions, we applied machine-learning-assisted boundary detection using CDeep3M,^27^ allowing high-throughput identification and quantification of microglial process architecture and neuronal interactions near Aβ plaques. In 5XFAD mice, microglia processes were retracted from neurons and Aβ plaques, whereas in 5XFAD:*Fgg^γ390–396A^* mice microglia extended processes into neurons and Aβ plaques (Figure 2C-D, Supplementary Videos 2, 3). Together, these data show that genetic inhibition of fibrin inflammatory domain signaling enhances microglial brain surveillance and restores microglia-neuron contacts in AD mice.

**Figure 2.**
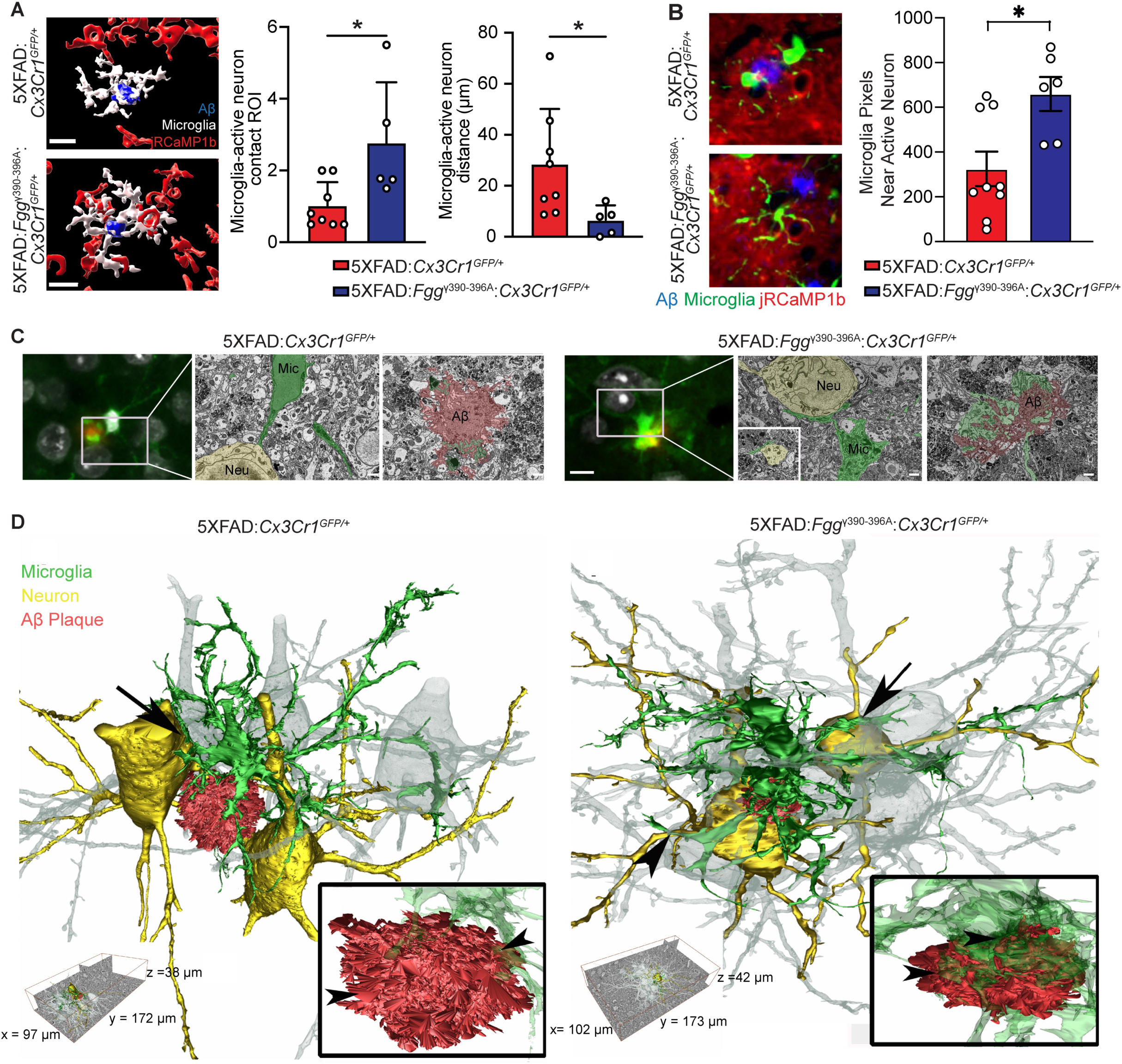
Blockade of the fibrin inflammatory domain restores microglia contacts with neurons in AD mice. (A) Microglia-neuron contact analysis. 3D reconstruction of microglia (GFP), active neurons (jRCaMP1b) and Aβ plaque (methoxy-X04) by Imaris and quantification of microglia-neuron contacts and distance of plaque-associated microglia to nearby active neuron. Data represents mean ± s.e.m. n = 8 5XFAD:*Cx3cr1*^GFP/+^ and n = 5 5XFAD:*Fgg^γ390-396A^*:*Cx3cr1*^GFP/+^ mice. \**P* < 0.05 by unpaired two-sample t-test. (B) Microglia response to spontaneous neuronal activity in awake mice. Average microglia voxels around each active neuron near Aβ plaque (within 50 μm). Data represents mean ± s.e.m. n = 9 5XFAD:*Cx3cr1*^GFP/+^ and n = 6 5XFAD: *Fgg^γ390-396A^*:*Cx3cr1*^GFP/+^ mice. \**P* < 0.05 by unpaired two-sample t-test. (C-D) SBEM (C) and 3D reconstruction (D) of microglial cell contacts with surrounding neurons and Aβ in 5XFAD and 5XFAD:*Fgg^γ390-396A^* mice. (C): Representative CLEM of microglia-neuron interactions in 5XFAD and 5XFAD:*Fgg^γ390-396A^* mice. Confocal images (left) show GFP-labeled microglia (green) and surrounding neurons labeled with DRAQ5 (white) near Aβ plaques labeled with methoxy-X04 (red). Corresponding SBEM image planes (right panels) illustrate ultrastructural details of microglial (micro) interactions with neurons (Neu) and Aβ plaques (Aβ) in each condition. Scale bars: 10 μm for confocal images, 1 μm for electron micrographs. (D) 3D reconstruction of microglia-neuron interactions within the neuronal connectome. Whole-body segmentation of microglia (green) and Aβ plaques (red) was performed using semi-automatic segmentation using CDeep3M, a machine-learning algorithm for large-scale EM data. Additionally, all neurons within the SBEM volume of each condition that physically interact with microglia were fully segmented (movie 2,3), including two representative neurons (yellow) and adjacent neurons (grey). Insets provide zoomed-in views of Aβ plaques and associated microglial fine processes, highlighting differences in microglial morphology and engagement between genotypes. Scale: 5XFAD, x = 97 µm, y = 172 µm, z = 38 µm. 5XFAD: *Fgg^γ390-396A^*, x = 102 µm, y = 173 µm, z = 42 µm.

### Blockade of the fibrin inflammatory domain restores the microglia-neuron glutamate and innate immune receptor crosstalk in AD mice

Neuroimmune interactions have emerged as key players in AD pathogenesis.^59^ We previously showed by single-cell RNA-sequencing (scRNA-seq) that microglia from 5XFAD:*Fgg*^γ390-396A^ mice have reduced oxidative stress and neurodegenerative gene expression, along with restoration of homeostatic signatures than microglia from 5XFAD mice.^45^ To determine the molecular mechanisms underlying microglia-neuron interactions, we performed single-nuclei RNA-sequencing (snRNA-seq) on the brains of 5XFAD and 5XFAD:*Fgg^γ390-396A^* mice and their littermate control WT and *Fgg^γ390-396A^* mice. A total of 165,252 nuclei from 14 mice were used for downstream analyses, and clustering analysis with uniform manifold approximation and projection (UMAP) identified 21 distinct clusters across all samples based on expression of known cell-type-specific gene makers, including excitatory neurons (L2/3 IT, L5 ET, L5 IT, L5/6 NP, L6 CT, L6 IT, L6 IT Car3 and L6b clusters), interneurons (Pvalb, Sst, Sst Chodl, Sncg, Vip, Lamp5), oligodendrocytes (oligo cluster), oligodendrocyte precursor cells (OPC cluster), astrocytes (Astro clusters), microglia and perivascular macrophage (Micro-PVM cluster), endothelial cells (Endo cluster), pericytes (Peri cluster), vascular and leptomeningeal cells (VLMC cluster) (Figure S3A).

Genetic blockade of fibrin inflammatory domain reduced aberrant transcriptional changes across multiple clusters in 5XFAD mice, including excitatory neuron clusters L2/3 IT (e.g., *Plag7, Bcas1, Adamts10, Dhcr24*), L5/6 NP (e.g., *Atm, Sema6d, Camk2g, Peg3*), L6b (e.g., *Slc8a3, Col19a1, Msi2, Cntn6*) and inhibitory neuron clusters Sst (e.g., *Aldh4a1, Grm3, Cdh13, Kcnip4*) and Pvalb (e.g., *Igsf1, Peg3, Gabra5*) (Figure 3A, S3B-C, Table S1). Gene set enrichment analysis (GSEA) showed that 5XFAD:*Fgg^γ390-396A^* mice exhibited restoration of AD-associated dysfunction in sterol biosynthesis, GPCR signaling, and glutamate import pathways in excitatory neurons, as well as hormone activity, ATP production, and metabolic pathways in inhibitory neurons (Figure 3B, Table S2). Consistent with previous reports,^60,61^ the microglial, astrocytic, oligodendrocytic, and neuronal clusters also displayed distinct 5XFAD-induced gene signatures than WT mice (Figure S3B, Table S1). In 5XFAD:*Fgg^γ390-396A^*mice, we found a broad rescue of AD-induced transcriptional signatures in non-neuronal cell types including astrocyte (e.g., *Cd9, Clu, Aqp4, Ifgbp5*), oligodendrocyte (e.g., *Stat3, Aldh1l2, Eya3, Abca1*), and endothelial cell (e.g., *Notum, Sp100, Nampt*) clusters (Figure S3C, S4A, Table S1). Gene Set Enrichment Analysis identified reduced microtubule motor activity in astrocytes, increased potassium transport and neurotransmitter secretion in endothelial cells and enhanced synapse assembly and neuropeptide signaling pathways in oligodendrocytes (Figure S4B, Table S2). Our findings suggest that inhibition of the fibrin inflammatory domain reduces AD signature genes across CNS innate immune, vascular, glia cells and neurons.

**Figure 3.**
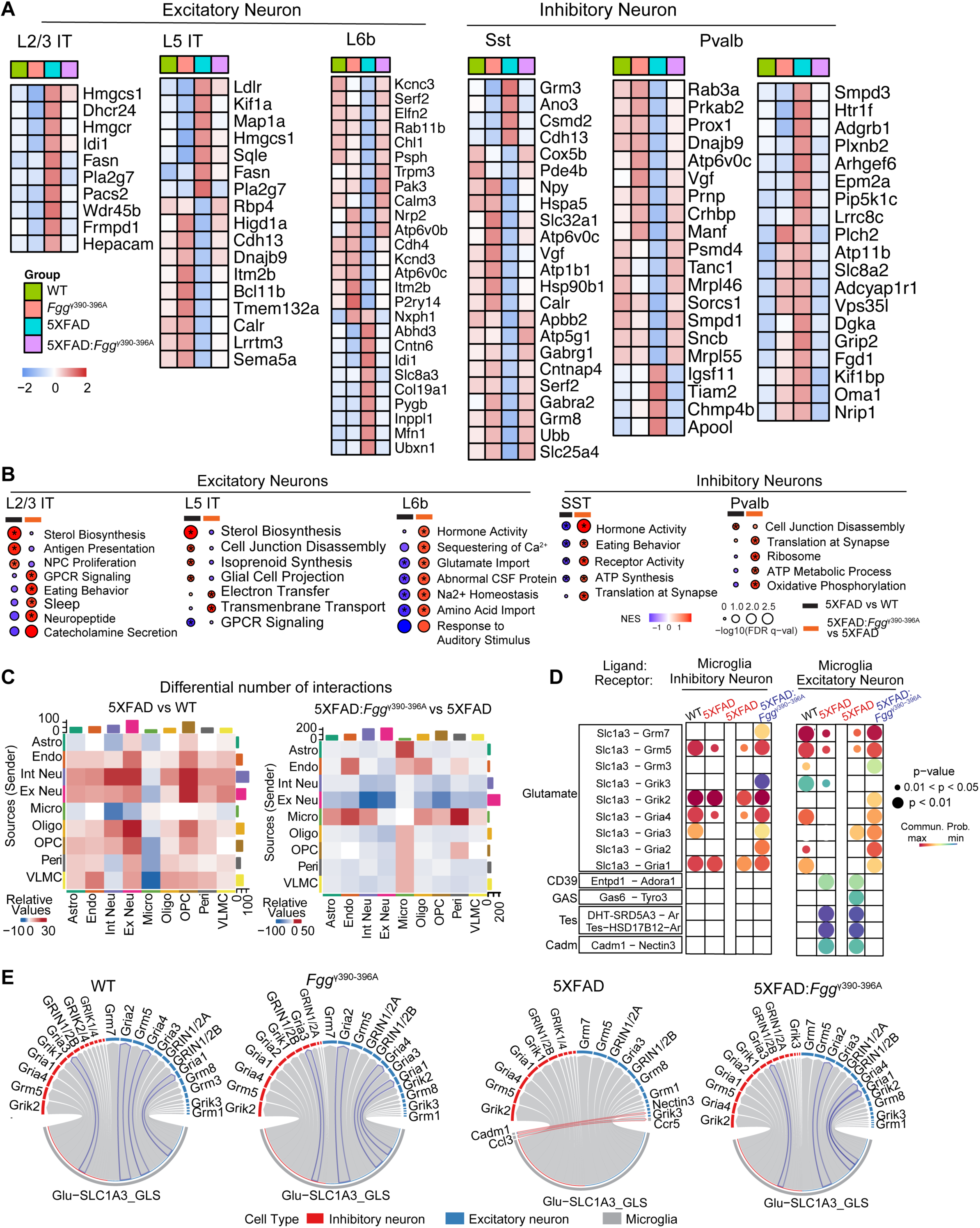
Fibrin disrupts the microglia-neuron interactome in AD mice. (A) Heatmap of significant DEGs in the excitatory neurons (L2/3 IT, L5 IT, L6b) and inhibitory neurons (Sst, Pvalb) mice from WT, *Fgg^γ390-396A^*, 5XFAD, and 5XFAD:*Fgg^γ390-396A^* mice by snRNA-seq (*P* < 0.05 in Pseudobulk analysis by 5XFAD vs WT and 5XFAD:*Fgg^γ390-396A^* vs WT). Colors indicate the average scaled expression level for each genotype. Data points are averages of *n* = 3 WT, n = 3 *Fgg^γ390-396A^*, n = 4 5XFAD and *n* = 4 5XFAD:*Fgg^γ390–396A^* mice. **(B)** Gene Set Enrichment Analysis (GSEA) of snRNA-seq data comparing 5XFAD vs WT and 5XFAD:*Fgg^γ390–396A^* vs 5XFAD in neuronal clusters using gene list ranked by log2FC from Pseudobulk analysis. (**C**) Cellchat analysis showing numbers of cell-cell interactions in 5XFAD versus WT (up) or 5XFAD*:Fgg^g390-396A^* versus 5XFAD (down) mice from snRNA-seq in combination of scRNA-seq microglia analysis (Mendiola et al 2023). Right bar plot depicts number of interactions from sender cell type cluster. Top bar plot depicts number of interactions received by target cell type cluster. (**D-E**) Cellchat analysis showing ligand-receptor changes between microglia and neurons (excitatory and inhibitory) in WT, *Fgg^γ390-396A^*, 5XFAD and 5XFAD:*Fgg^γ390–396A^* mice. (**D**) Dot plots showing significantly altered pathways and ligand receptor pairs. (**E**) Circle plots showing significantly altered ligand-receptor pairs sent from microglia to excitatory neurons (blue) and inhibitory neurons (red).

To identify the molecular mediators underlying microglia-neuron interactions governed by fibrin inflammatory signaling, we determined the ligand-receptor interactions and cell-cell communication networks from snRNA-seq and scRNA-seq datasets using Cellchat. snRNA-seq data were used for all neuronal and glia cell types. Due to the low abundance of the microglia populations in the snRNA-seq dataset, we used scRNAseq data from FACS-sorted microglia were incorporated to improve microglial transcriptomic coverage.^45^ We first computed a cell-cell interaction map across all cell types to identify the number of predicted dysregulated pathway interactions. 5XFAD mice exhibited a marked reduction in microglial interactions with astrocytes, inhibitory and excitatory neurons, oligodendrocytes, and the endothelial, OPC, VLMC, and pericyte clusters, many of which were reversed in 5XFAD:*Fgg^γ390-396A^* brains (Figure 3C, S4C). We next compared the changes of specific ligand-receptor pair pathways in microglia (sender) and neurons (receiver). In 5XFAD mice, we found a reduction of glutamate pathway pairs between microglia and both excitatory and inhibitory neurons, while CD39-Adora, GAS6-Tyro3, Testosterone, and Cadm-Nectin3 pathway pairs were increased between microglia and excitatory neurons; all of these pairs were normalized in 5XFAD:*Fgg^γ390-396A^* mice (Figure 3D and 3E, Table S3). These data suggest that fibrin inflammatory signaling impairs the communication of microglia with excitatory and inhibitory interneurons by disrupting key molecular pathways involved in neuronal network synchronization.

### CSF fibrinogen positively correlates with vascular, immune and synaptic biomarkers in AD patients

Since we identified a causal role for fibrin in driving expression of inflammatory and synaptic dysfunction genes in AD mice, we tested whether these biological processes correlate with fibrinogen levels in the CSF of AD patients in independent analysis of human samples. We performed unbiased proteomic profiling of CSF from 57 AD patients (ages 51–81; both male and female) using the SomaLogic 7K platform (Figure 4A). A total of 7260 proteins were robustly detected and included in downstream correlation analyses (Table S4). Fibrinogen and D-dimer levels measured by SomaLogic were highly correlated with fibrinogen concentration in the CSF obtained by ELISA (Supplementary Figure 5, Table S5), validating the Somascan aptamers. In the CSF of AD patients, 296 proteins positively correlated with CSF fibrinogen, while 94 proteins were negatively correlated (Figure 4B, Table S5). By pathway analysis, we found that CSF fibrinogen positively correlated with complement activation, coagulation, inflammatory response and cholesterol homeostasis, but fibrinogen negatively correlated with Peptidyl−threonine phosphorylation and the APP catabolic process (Figure 4C). Cell type enrichment analysis showed that fibrinogen-associated proteins in the CSF largely originated from glutamatergic neurons, GABAergic neurons, and endothelial cells (Figure 4D). CSF fibrinogen levels positively correlated with markers of BBB dysfunction and coagulation (HABP2, F11, PLG, F13B, F10, F9, F7, KNG1, TFPI, KLKB1), inflammation (C3, C5, C6, C9, CR1, CFP, IL1R2, IL1RAP, IL19, IL16, LCN2, HEXB), and neuron and synaptic markers (NRG1, NXPH1, SHH, ITM2B, MAP2K4) (Figure 4E). Overall, human AD biomarker data reveal a correlation of CSF fibrinogen with vascular, immune and neurodegenerative biomarkers in AD patients.

**Figure 4.**
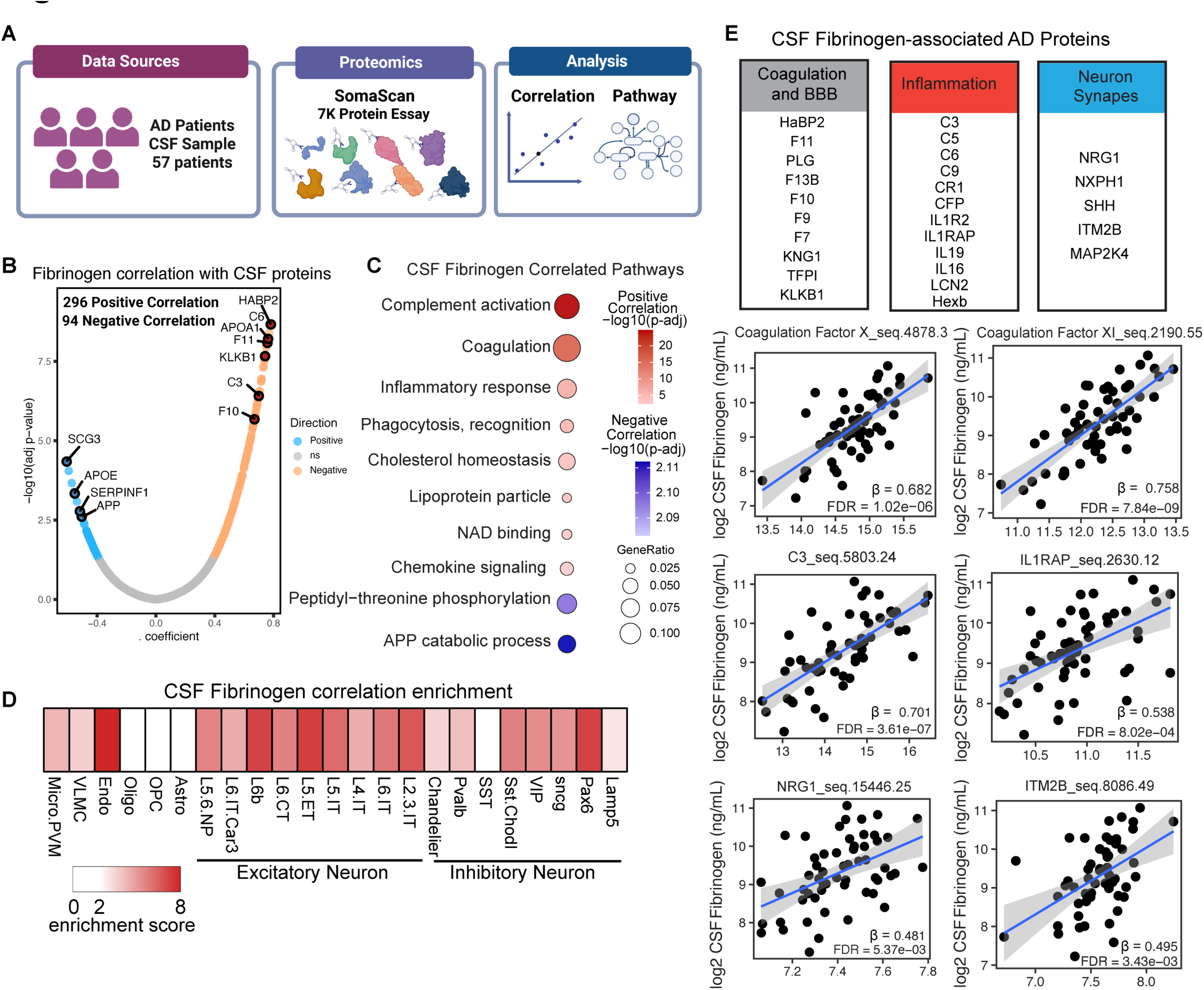
Fibrin correlates with vascular, immune and synaptic dysfunction biomarkers in the CSF of AD patients. (**A**) Schematic of unbiased analysis of human AD CSF samples using Somascan. (**B**) Volcano plots of CSF proteins significantly correlated with CSF fibrinogen (adjusted *P* < 0.05 by Pearson correlation). (**C**) Gene ontology pathway analysis of CSF proteins positively and negatively associated with fibrinogen in human AD patients. (**D**) Fibrinogen correlation enrichment score of each brain cell type calculated by overlaying significantly correlated analytes with the Human M1 10x database (https://brain-map.org/). (**E**) Up, top markers of coagulation and BBB, inflammation, and synapse significantly correlated with CSF fibrinogen in AD patients adjusted P < 0.05 by Pearson correlation. Bottom, representative dot plots showing the correlation between CSF fibrinogen and markers of BBB, inflammation, and synapse.

## Discussion

Our study identifies an extrinsic, blood-derived mechanism that induces microglia-mediated neuronal hyperactivity, with important implications for neuronal network dysfunction in AD. We demonstrate that fibrin inflammatory signaling impairs microglial brain surveillance, disrupts microglia-neuron contacts and promotes aberrant neuronal activity. In AD mice, genetic disruption of the fibrin inflammatory epitope improves microglial brain surveillance and restores neuronal activity to physiological levels, consistent with reduction of neurodegenerative and inflammatory gene expression in microglia.^34,45^ Restoration of transcriptional signatures across excitatory and inhibitory neurons, astrocytes, and endothelial cells suggests an apical role for fibrin-induced microglial activation driving pathogenic remodeling of the neurovascular unit in AD that leading to neuronal hyperactivity. By elucidating the fine architecture of the dynamic microglial-neuron contacts around Aβ plaques at the ultrastructural level, we demonstrate that inhibition of the fibrin inflammatory domain can restore the microglia-neuron interactome even in the presence of Aβ plaques. Our study identifies vascular signals as potent Aβ-independent drivers of microglia dysfunction and neuronal hyperactivity. These results are consistent with CSF fibrinogen as an early amyloid-independent neurodegeneration biomarker.^42^ By constructing a ligand-receptor atlas of microglial interactions with inhibitory and excitatory neurons, we reveal an unanticipated role for fibrin in disrupting the microglia-neuron molecular interactome, including glutamatergic and innate immune signaling pathways in AD. Given that vascular dysfunction and neural network abnormalities are early hallmarks of AD, our data suggest that BBB disruption and cerebrovascular pathology may alter the rules of neuron-microglia communication that underlie brain hypersynchrony. Furthermore, 5XFAD:*Fgg^γ390-396A^* mice have reduced hyperexcitability shown by chronic EEG recordings.^62^ As neural hyperactivity and hyperexcitability are observed at early stages of disease often preceding profound neuronal loss,^63–65^ our findings suggest that fibrin is an amyloid-independent regulator of immune-mediated circuit dysfunction. By identifying fibrin as an apical signal disrupting the molecular connections between microglia and neurons leading to hyperactivity, our study provides a molecular mechanism for cerebrovascular pathology as an early driver of neural network dysfunction. Together, these findings support targeting fibrin inflammatory domain as a promising therapeutic intervention for the restoration of neuronal networks in AD.

Microglia sense neuronal activity and regulate network excitability via pathways including G-protein coupled receptors, CD39/Adora and ATP/P2Y12.^25–27,66^ Our data show that genetic blockade of fibrin inflammatory domain restored microglia-neuron communication molecular pathways related to seizures and AD including SLC1A3, GAS6/TAM and CD39/Adora signaling pathways. SLC1A3, which encodes the glutamate transporter EAAT1, is expressed in activated microglia and astrocytes,^67^ and mutations in SLC1A3 are associated with neurological disorders such as episodic ataxia, hemiplegia and seizures,^68^ as well as increased risk of epileptogenesis following traumatic brain injury.^69^ Since fibrin-induced microglial activation suppressed microglial SLC1A3-neuronal glutamate ligand-receptor interactions, fibrin may promote neuronal hyperactivity by impairing glutamate sensing by microglia. Additionally, genetic inhibition of fibrin-CD11b signaling restored GAS6-Tyro3 interactions. GAS6, a potent agonist of TAM receptors (Tyro3, Axl, MerTK), regulates Aβ plaque clearance^70–72^ and is associated with thrombosis, worse behavioral outcomes in AD mice and impaired remyelination in multiple sclerosis patients.^73^ Moreover, fibrin-induced microglial activation was required for enhanced CD39/Adora G-coupled receptor signaling in AD mice, which is involved in microglia-mediated regulation of network hyperexcitability.^25^ Together, our data suggest cross-talk between fibrin integrin signaling and G protein-coupled and TAM receptors in microglia, consistent with prothrombotic platelet signaling transduction orchestrated by P2Y12R, A2R, TAM and fibrin α_IIb_β_3_ integrin receptor activation.^74^ Fibrin-induced activation of integrin receptors may serve to integrate cerebrovascular dysfunction and coagulation with associated microglia responses required for network synchronization, as well as to coordinate signal transduction pathways and cell-cell interactions involved in BBB integrity, scar formation and neurodegeneration.

We identified novel neurodegeneration-associated proteins that correlate with CSF fibrinogen levels in patients with AD, including markers of BBB disruption, coagulation, neuroinflammation, and synapse loss. These findings are consistent with reports that the CSF/plasma fibrinogen ratio correlates with cognitive impairment in 2,171 AD patients^13^ and that CSF fibrinogen levels in cognitively normal older adults correlate with cognitive decline, inflammatory markers, synaptic injury, and vascular dysfunction independent of amyloid pathology.^42^ In our study, CSF from AD patients from the UCSF cohort, fibrinogen strongly correlated with multiple coagulation factors and complement components. Supporting the relevance of this observation, fibrin induces coagulation factor X expression in neurotoxic microglia and anticoagulants protect from neuroinflammation and cognitive impairment in AD models.^75,76^ Complement is also elevated in CSF from AD patients^77^ and correlates with cognitive deficits,^13,78^ and pharmacologic inhibition of the fibrin inflammatory domain by antibody 5B8 decreases complement pathways in AD mice.^43^ Notably, CSF fibrinogen was significantly correlated with many AD risk genes (*APOE*, *CR1*, *IL1RAP*, *APP*, *CST3*), further supporting its relevance to AD pathophysiology. In addition, CSF fibrinogen positively correlated with neuronal and synaptic markers, including NRG1, a presynaptic protein which is increased in AD patients,^79^ consistent with an earlier finding that fibrinogen levels correlate with synaptic vulnerability during normal aging.^42^ Together with our experimental evidence demonstrating a causal role for fibrin in regulating inflammation and neuronal activity in mice, AD biomarker studies position fibrin-induced microglia activation as a central mechanism linking vascular pathology, neuroinflammation and neuronal dysfunction in AD and related dementias.

By developing a cost–benefit conflict test, we identified fibrin inflammatory signaling as a driver of increased high-risk behavior in AD mice. This is in line with high-risk-taking behaviors in early-stage AD patients indicative of executive dysfunction independent of memory impairment.^54,80^ Our study suggests that targeting fibrin may improve early behavioral alterations associated with executive dysfunction preceding memory impairment. Identification of fibrin inflmmatory signaling as a driver of high-risk choice bevavior suggests a central role for fibrin in immune-mediated cognitive impairment, as also shown by the protection of 5XFAD:*Fgg^γ390-396A^* mice from deficits in learning and memory, spontaneous behavioral abnormalities, anxiety, hyperactivity, epileptiform activity and sleep disruptions.^34,47,62^ Since decision-making depends on prefrontal cortex circuits that are impaired in AD patients,^81–84^ our findings suggest a role for vascular and immune mechanisms regulating neuronal circuits controlling risk-taking behaviors. These findings are unlikely due to retinal dysfunction, as the 5XFAD-C57B/6 line tested in this study does not carry the retinal degeneration allele *Pde6b^rd1^* and has normal visual acuity at 8 months of age.^85^ Together, these findings suggest that fibrin-induced microglial activation is required for neuronal dysfunction and a broad range of cognitive deficits associated with AD and related dementias.

Our study has several limitations. Although we identified novel fibrin-mediated ligand-receptor pairs involved in microglia-neuron interactions in AD mice and uncovered CSF proteins correlated with fibrinogen levels in AD patients, future functional validation studies will be required to test the causal contributions of these downstream fibrin-triggered neuroimmune pathways to neural hyperactivity in AD. Additionally, although excitation-inhibition imbalance and network hypersynchrony are proposed as early drivers of cognitive dysfunction in AD,^3,4,20^ it remains unclear whether microglia respond equally to excitatory and inhibitory neurons or whether fibrin differentially regulates excitatory and inhibitory neuronal activity. Since blocking the fibrin inflammatory domain reduces AD-related astrocyte gene expression, future studies will determine the relative roles of microglia and astrocytes in the regulation of fibrin-induced neuronal hyperactivity. Our findings were generated in the 5XFAD model of AD. Fibrin plays a role in several AD models—including 5XFAD, APP/PS1, TgCRND8, PDAPP, Tg2576, the non-overexpressing rat Familial Danish Dementia (FDD)-KI, APP^SAA^-KI, APP^NLG^-KI mice, and the TgSwDI mouse model with robust CAA pathology.^33–36,86^ Consistent with this study and our prior work in 5XFAD mice,^34,43,45^ fibrin depletion protects from microglia activation in multiple strains of AD models, including the TgSwDI and TgCRND8 mice,^35,87,88^ suggesting that fibrin is a general mechanism of microglia activation in amyloid-driven neurodegeneration. In addition to protection from high-risk behavior in AD mice (this study), fibrin blockade exerts protection in a broad range of cognitive behavioral tests assessing learning and memory, hyperactivity and spontaneous behavior across different AD mouse models. ^34,44,47,62,87,88^ Both male and female mice were used for the experiments, but the study was not powered to evaluate sex-specific effects. Our assay to assess risk-taking behavior does not exclude potential confounding effects of learning and memory or sex differences on decision making. Future studies with additional decision-making and reward-seeking behavioral tests will be required to evaluate potential effects of learning and memory and sex in risk-taking behaviors, as well as across different AD mouse models with vascular pathology and neuroinflammation.

Together, data from biomarkers and neuropathology, as well as genetic-loss-of function studies establish the fibrin inflammatory domain as a key driver of immune-mediated neurodegeneration and cognitive impairment in AD and related dementias.^32–34,45,47^ The antibody 5B8, which targets the fibrin γ_377–395_ inflammatory domain, protects from autoimmune, infectious and amyloid-driven neurodegeneration without affecting hemostasis.^43,89,90^ In 5XFAD mice, therapeutic administration of 5B8 reduces microglia activation, inflammatory response, and complement signaling with neuroprotective effects.^43,45,90^ THN391, a humanized high affinity variant of 5B8, has completed Phase 1 safety trials and was safe and well tolerated with no impact on hemostasis, and has now advanced in Phase 1b trials in patients with AD (clinical trials.gov: NCT06814730).^86,89^ Pharmacologic targeting of the fibrin γ_377–395_ inflammatory domain by using 5B8 or THN392, a variant of THN391 with a murine Fc domain enabling chronic mouse dosing, reduced epileptic phenotype in 5XFAD mice. ^62^ Cerebrovascular dysfunction associated with high fibrinogen CSF levels has emerged as an early and amyloid-independent pathway associated with inflammation, synaptic degeneration, tau pathology, and cognitive dysfunction in AD and aging.^2,6,13,38,40,42,91^ The efficacy and safety targeting the fibrin γ_377–395_ inflammatory domain suggests that fibrin-immunotherapy could be a therapeutic strategy to selectively suppress neurotoxic inflammation and protect from neurodegeneration, neuronal dysfunction and network hyperexcitability in AD. Fibrin targeting therapies may be used alone or in combination with Aβ-lowering immunotherapies to improve cognitive function in AD. Since fibrin is present in the amyloid-related imaging abnormalities (ARIA),^41^ fibrin immunotherapy may also reduce the adverse effects of microhemorrhages and concomitant neuroinflammation of amyloid-lowering therapies.^86,89^ Together, these findings underscore fibrin-targeting antibodies as a promising therapeutic strategy for vascular pathology, pathogenic immune activation, and neuronal dysfunction not addressed by other treatments.

## RESOURCE AVAILABILITY

### Lead contact

Requests for further information and resources should be directed to and will be fulfilled by the Lead Contact, Katerina Akassoglou

### Material availability

Materials will be made available upon request under the appropriate material transfer agreements.

### Data and code availability

snRNAseq datasets will be publicly available upon publication. The microglia scRNA-seq datasets used for the Cellchat analysis reported in Mendiola et al 2023 (Ref. 48) are deposited in the Genome Expression Omnibus under SuperSeries accession number GSE229376.

## ACKNOWLEDGEMENTS

We thank the Histology and Light Microscopy Core of Gladstone Institutes and Katy Claiborne for editing. The Gladstone FACS Core acknowledges US National Institutes of Health (NIH) grant S10 RR028962 and the James B. Pendleton Charitable Trust. Dr. Miller receives grants P0544014 for the UCSF FTD Core from the Bluefield Project to Cure FTD and R01AG057234 from the NIH/NIA for the US-South American Initiative for Genetic-Neural-Behavioral Interactions in Human Neurodegenerative Diseases. This research was supported by Warren Alpert Distinguished Scholar Award (Z.Y.), Brightfocus Postdoc Fellowship Award A2021019F (Z.Y.); Kaganov Scholarship for Excellence in Neuroscience (Z.Y.); NIH/NINDS K99 NS126707 (A.S.M.); NIH grants U24 NS120055, R24 GM137200, R01 GM138780 and S10 OD021784 (M.H.E.); RF1 AG062234 (J.J.P.), R01 AG062629 (J.J.P.), and R01 AG092683 (J.J.P.); National Science Foundation NSF2014862-UTA20-000890 (M.H.E.); philanthropic gifts from Edward and Pearl Fein, Robert Hamwee, the Dolby Family Fund, the Spangler Foundation and the Simon Family Trust (K.A.); the Brightfocus Award A2025013S (K.A.), the Alzheimer’s Association Zenith Award 26-1447450 (K.A.), NIH grants NIA RF1 AG064926 (K.A., J.J.P., M.H.E.), NINDS R35 NS097976 (K.A.) and R35 NS143067 (K.A.).

## AUTHOR CONTRIBUTIONS

Conceptualization: KA, ZY, ASM, KL, KYK, YY, FME, MHE, JJP. Data curation: Z.Y., A.S.M., K.Y.K., J.N.K., F.M.E, M.H.E, K.A.; Designed Experiment: ZY, ASM, KYK, EAB, YY, FME, MHE, JJP, KA. Performed Research: ZY, ASM, KYK, YY, KLeng, EAB, AMF, MFV, DCB, MM, RTognatta, MPSA, RA, BC, KNP, KH, JKR. Formal Analysis: ZY, ASM, KYK, YY, KLeng, RS, AA, MT, NE, RT, JNK, NK, MFV, DCB, MM, BG, KH, KA, MHE, F.M.E. Data Interpretation: ZY, ASM, KL, KYK, YY, KLeng, RS, AA, MT, NE, RT, JNK, NK, RTognatta, MFV, DCB, MM, BG, KH, F.M.E., MHE, JJP, KA, Software: KLeng, RS. Visualization: ZY, ASM, KYK, MT, LK. Writing-Original Manuscript: KA, ZY, ASM, Writing: Review & Editing: All Authors. Supervision: KA, MHE, FME., JJP. Funding Acquisition: KA, JJP, MHE, FME, ZY, ASM. Project Administration: KA.

## STAR★METHODS

### Key resources table

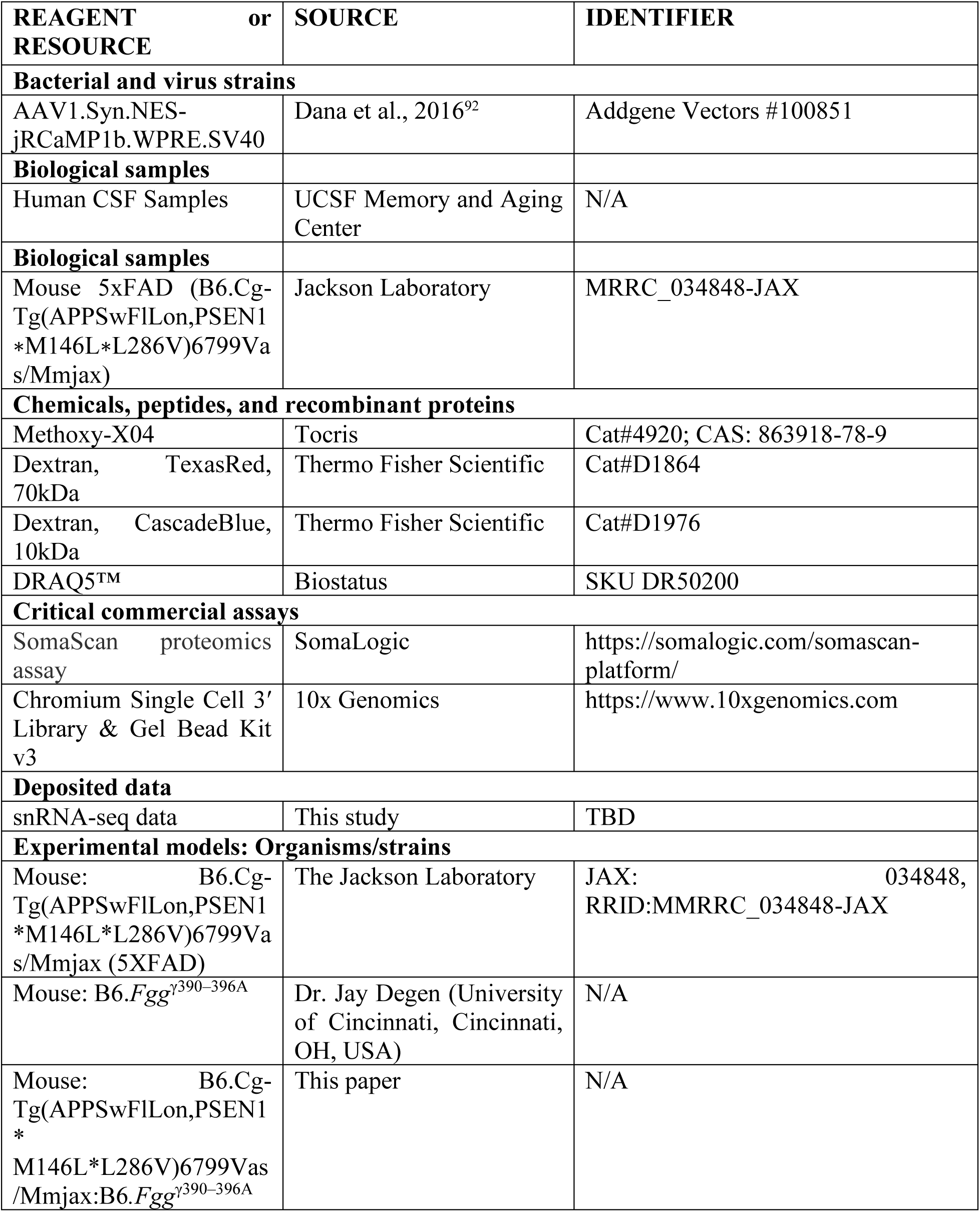
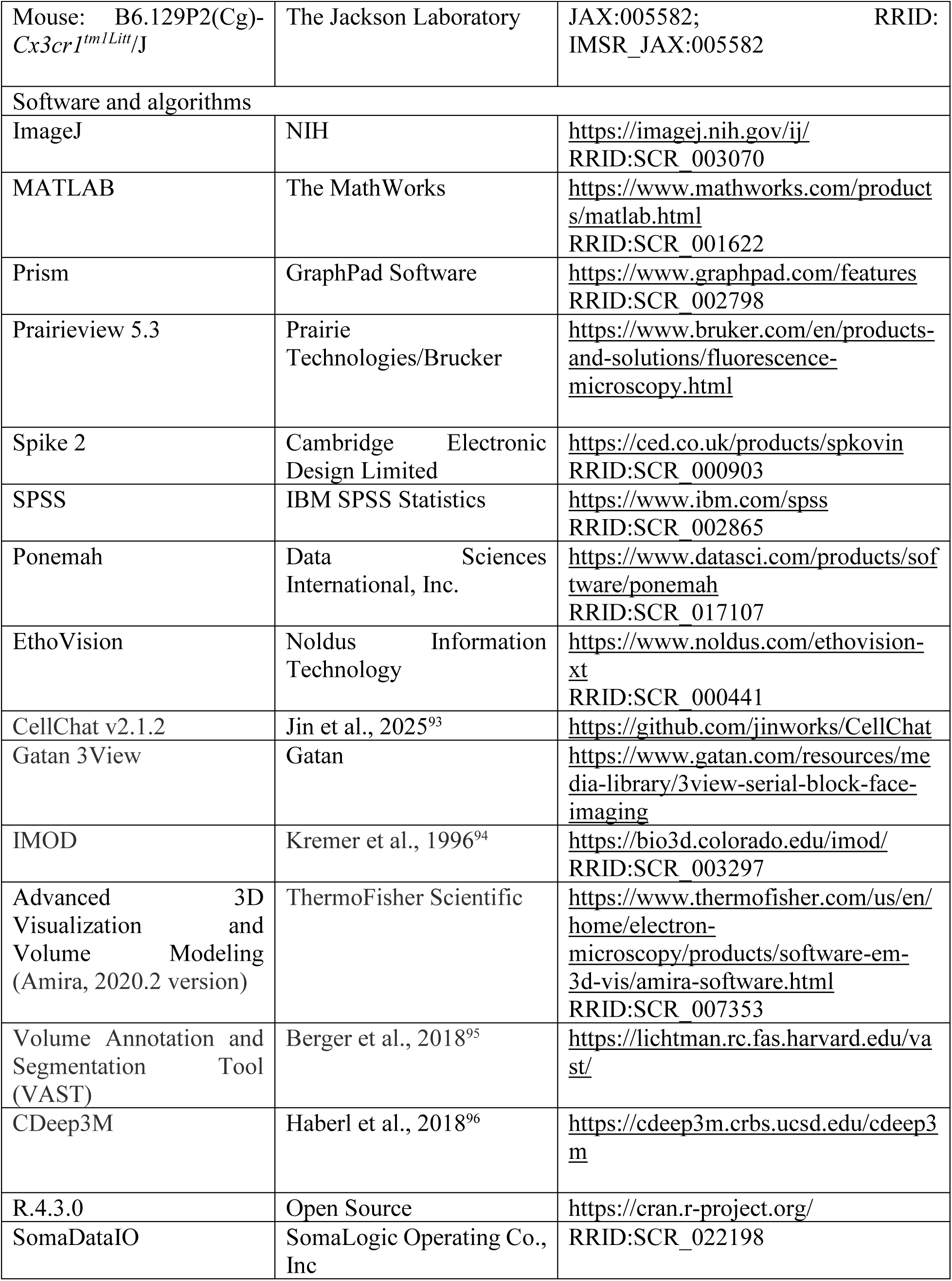
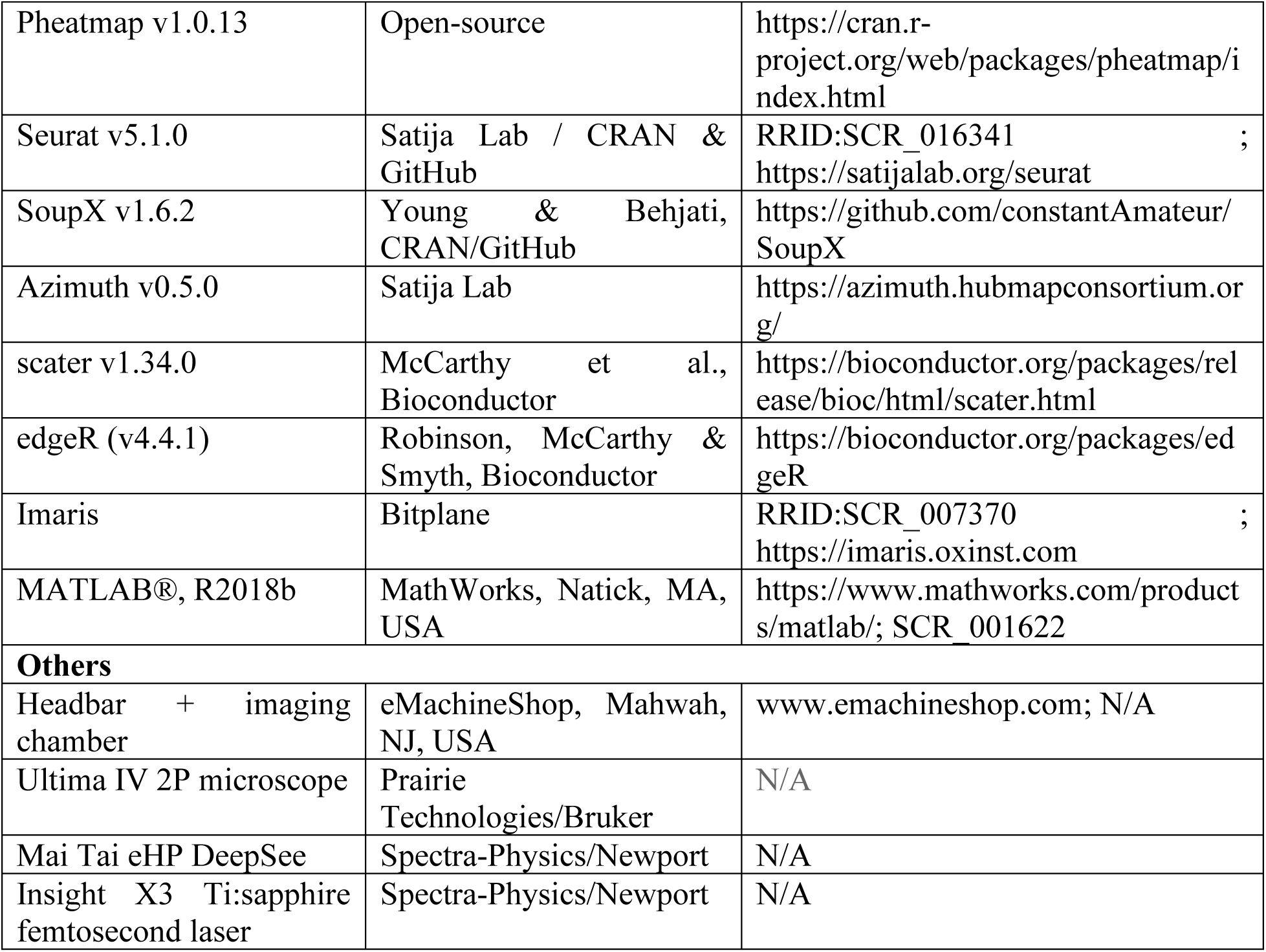

### EXPERIMENTAL MODEL AND STUDY PARTICIPANT DETAILS

#### STUDY PARTICIPANTS

Participants were drawn cross-sectionally from ongoing longitudinal studies at the UCSF Memory and Aging Center and included individuals with biomarker-positive Alzheimer’s disease based on cerebrospinal fluid A/T/N or positive amyloid status on positron emission tomography (PET) obtained within 12 months of the biospecimen collection. Written informed consent was obtained from all participants, and all procedures were approved by the UCSF Committee on Human Research and conducted in accordance with the principles of the Declaration of Helsinki.

#### EXPERIMENTAL MODELS

B6.129P2(Cg)-Cx3cr1tm1Litt/J(*Cx3cr1^GFP/GFP^*) and B6.Cg-Tg(APPSwFlLon, PSEN1*M146L*L286V)6799Vas/Mmjax(5XFAD) mice were obtained from the Jackson Laboratory. *Fgg^γ390–396A^* mice were obtained from Dr. Jay Degen (University of Cincinnati, OH, USA). The 5XFAD line was maintained by crossing male 5XFAD mice with female C57BL/6 mice. The 5XFAD:*Fgg^γ390–396A^* mice were generated by crossing male 5XFAD mice with *Fgg^γ390–396A^* mice and were maintained by crossing male 5XFAD:*Fgg^γ390–396A^* mice with female *Fgg^γ390–396A^*mice. The 5XFAD:*Cx3cr1^GFP/+^* mice were generated by crossing male 5XFAD mice with female *Cx3cr1^GFP/GFP^* mice. The 5XFAD:*Fgg^γ390–396A^* :*Cx3cr1^GFP/+^* mice were generated by crossing male 5XFAD mice with female *Fgg^γ390–396A^* :*Cx3cr1^GFP/GFP^* mice. Mice were housed under a 12:12 light/dark cycle at 55% ± 5% relative humidity and at a temperature of 20 ± 2°C with ad libitum access to standard laboratory chow and water ad libitum. Mice that had undergone cranial window surgery were housed individually to prevent damage to the cranial window. Mouse ages are indicated for each experimental procedure. All animal experiments were performed following the guidelines set by the Institutional Animal Care and Use Committee at the University of California, San Francisco.

### METHOD DETAILS

#### Cranial window surgery and stereotaxic injections

The craniotomy window surgery was performed as previously described with minor modifications.^34,97^ Mice were anesthetized with intraperitoneal (i.p.) ketamine and xylazine. The head was shaved, disinfected with Betadine and 70% ethanol, and locally anesthetized with 2% Lidocaine (subcutaneous, s.c.). Subcutaneous connective tissues were removed using forceps to expose the skull. Once the injection site was identified, the bone was thinned using a drill, followed by a stereotaxic injection of 1 µl of pAAV.Syn.NES-jRCaMP1b.WPRE.SV40 (Addgene, 100851) into the somatosensory cortex (AP = −1.7 mm, ML = 2.65 mm, DV = −0.85 mm, relative to bregma) using a 33G beveled Hamilton syringe at a speed of 60 nL/min. Two self-tapping bone screws were placed on the skull to anchor the dental cement. A 4 mm circular outline was drawn using a drill, and the bone on the line was further thinned to create a circular groove. A custom-designed steel head bar with an imaging chamber (eMachineShop, Mahwah, NJ, USA) was affixed to the skull using cyanoacrylate glue and C&B Metabond® Quick Adhesive Cement (Parkell). The bone inside the imaging chamber was further cleaned and thinned with a drill, and the chamber was filled with pre-warmed artificial cerebrospinal fluid (ACSF). The skull within the circle was carefully removed and replaced with a 4 mm cover slip, which was sealed with Flow-it ALC (Pentron) and cured under UV light. All imaging experiments were conducted at least two weeks post-surgery.

#### *In vivo* 2P Microscopy

All *in vivo* imaging experiments were performed using an Ultima IV 2P microscope (Prairie Technologies/Bruker) equipped with a Mai Tai eHP DeepSee and an Insight X3 Ti:sapphire femtosecond laser (pulse width <120 fs, tuning range: 690–1,040 nm for Mai Tai, 680–1,300 nm for Insight X3, repetition rate: 80 MHz; Spectra-Physics/Newport). Microscopy, laser control, and image acquisition were managed using PrairieView software (v.5.4, Bruker). Imaging was performed at a depth of 100–200 µm below the dura to assess neuronal activity and microglial response. To prevent overheating, the maximum laser power was limited to 40 mW in all experiments. Depending on the resolution requirements, either a Nikon ×40, 0.8 NA or an Olympus ×25, 1.05 NA water-immersion objective lens was used. A GaAsP PMT (Hamamatsu, H7422PA-40) was employed for neuronal activity imaging, while a multi-alkali type PMT was used for other channels.

#### *In vivo* imaging of neuronal activity and microglial surveillance in awake mice

Eleven days post-surgery, mice were habituated to the imaging stage by being head-fixed for 10 minutes per day for three consecutive days. Methoxy-X04 was injected one day before imaging to label amyloid plaques. For each region of interest (ROI), a Z-stack (z-series) of 80–100 µm was first acquired in the jRCaMP1b and Methoxy-X04 channels at layer II/III (100 µm below the dura) using galvo scan mode (512 × 512 pixels, 1.5 Hz, ×2 optical zoom) with a ×25 objective (Olympus, 1.05 NA). Areas containing amyloid plaques, as indicated by Methoxy-X04 labeling, were selected for neuronal activity imaging. Spontaneous neuronal activity was recorded for 1 minute (T-series) using spiral scan mode (256 × 256 pixels, 12 Hz, ×2 optical zoom) with a ×25 objective (Olympus, 1.05 NA). For microglia imaging in awake mice, a Z-stack of 80–100 µm was acquired for the jRCaMP1b (1,100 nm), Methoxy-X04 (810 nm), and GFP (920 nm) channels at layer II/III (100 µm below the dura) using galvo scan mode (512 × 512 pixels, 1.5 Hz, ×2 optical zoom) with a ×25 objective (Olympus, 1.05 NA). To assess microglial responses to spontaneous neuronal activity, prior to the a Z-stack of 80–100 µm image, three-minute time-series images were acquired on a single plane using galvo scan mode (512 × 512 pixels, 1.5 Hz, ×2 optical zoom) with a ×25 objective (Olympus, 1.05 NA) for representative image and 1 minute (T-series) using spiral scan mode (256 × 256 pixels, 12 Hz, ×2 optical zoom) with a ×25 objective (Olympus, 1.05 NA) was taken for segmentation.

#### *In vivo* imaging of microglia in KX-anesthetized mice

Methoxy X04 was injected one day prior to imaging to label amyloid plaque. Two weeks post cranial window surgery, mice were anathesized with Ketamine and Xylazine and put on the imaging stage in a heat-regulated chamber. For microglia surveillance, a serial of a Z-stack (z-series) of 60-80 um image was acquired in the GFP (920nm) and Methoxy-X04 (810 nm) channel at layer II/III (100 below dura) using galvo scan mode at 512 × 512 pixels, 1.5 Hz and ×2 optical zoom with a ×25 objective (Olympus, 1.05 NA) for each ROI. The Z-stack was imaged every 2 min for 50 min. For microglia motility imaging, a serial of a Z-stack (z-series) of 25-30 um image was acquired in the GFP (920nm) and Methoxy-X04 (810 nm) channel at layer II/III (100 below dura) using galvo scan mode at 512 × 512 pixels, 1.5 Hz and ×3 optical zoom with a ×40 objective (Nikon, 0.8 NA) for each ROI. The Z-stack was imaged every 30 seconds for 15 min.

#### Cost-Benefit Conflict (CBC) Test

The CBC test was performed with modifications as previously described. ^56^ All mice with mixed genotypes were group-housed in their home cages. 40mL of Horizon Organic 1% Low Fat Chocolate Milk purchased from a local grocery store was put in the cage for mice. Mice were allowed to drink the chocolate milk without performing any task for 2 days. Mice first performed the forced-choice training. The T-maze apparatus (stem 30 x 10cm, arm length 30cm each) was used for the CBC test. Each maze arm was fitted with a food well (50mL test tube cap) filled with either pure (high reward) chocolate milk or diluted 80% chocolate milk (low reward). Two Lume Cube panels with 70% 3200K of power each were secured at the right arm of the T-maze. Prior to training, the mice were transferred from the holding room to the behavior testing room and habituated in the room for 1 h. Each mouse underwent 4 days of forced-choice training, 3 trials per day, 3 min per trial. During training session, the Lume Cube panels were turned on to overcome potential fear-induced response, as described.^56^ We paired the high reward with bright light (~1200 lumens, a high cost) and low reward with a dim light (~30 lumens, a low cost). One arm of the maze was blocked off so that the mice were forced to explore the only open arm. The open arm was alternated between each forced-choice training trial. The mice followed the same running order across trials. The latency to drink and behaviors including grooming, staying at the start location or open arm, and smelling without licking the reward were recorded by the experimenter to determine whether the mouse is well-trained by drinking the reward. If the mouse was able to consistently drink the reward (more than 2 out of 3 trials) in the last training day, the mouse would be tested in the following free-choice experimental trials. After forced-choice training, mice performed free-choice experimental trials. Mice were given 20 free-choice experimental trials, 5 trials per day. The set-up was identical to the forced-choice training session; however mice were released onto the stem of the T-maze with both arms of the maze open for exploration. A mouse was considered to have made a “decision” when it drank at lease some chocolate milk from the food well of an arm. No decision was recorded if a mouse chose not to drink from either food well after 3 min on the maze. If the mouse did not drink the reward within 3 min, the trial was considered as omission. The trial ended after the mouse drank some chocolate milk. The time each mouse took to make a “decision” (drink the reward from either food well) was recorded as the latency to choose. Different arm entries made by the mouse prior to the “decision” were also recorded. Data from the 5 free-choice experimental trials for each mouse each day were averaged to find percentages of decisions in which a mouse chose the high-risk/high-reward alternative over 4 continuous days. If there were >10 out of 20 omitted trials, the mouse would be excluded from the quantification of decisions and latency to choose. If there were > 2 out of 5 omitted trials in one day, but no more than 10 omissions in total, the decisions made by the mouse in that day and only that day would be excluded. For statistical analysis, our study was a 3 Genotype x 4 days of repeated measures design. We had 5 measures from the CBC test: percentages of high risk/high reward decisions, the latency to choose, number of omissions, light entry ratio, and average entries per trial. The areas under the curve of the percentages of high risk/high reward decisions were calculated for each group to determine the genotype effect by Kruskal-Wallis test and Holm-Sidak’s multiple comparisons test. We used a generalized linear mixed effects model (GLMM)^98^ to model the change in the mean log odds of mouse making a high risk/high benefit choice across trials on a given day, due to genotype, due to the day of the trial and interaction between these variables. The Mouse ID was used as a random effect in this model. The significance of genotype in the above model fit was based on an ANOVA test comparing the above model fit to the fit of the GLMM model including only the day of the trial. The differences in the mean log odds estimates between pairs of genotypes on a given day are estimated using the emmeans function implemented in the emmeans package. The p-values from each of these comparisons are corrected for multiple testing using the Holm procedure implemented in the p.adjust function in R.

#### Image Analysis for *in vivo* 2P imaging

The open-source package CaImAn was used to analyze spontaneous neuronal activity. Unless otherwise specified, the default settings were used, with the following parameters adjusted for analysis: frame rate = 12.5 Hz, decay time = 0.6 seconds, and gSig = np.array([4, 4]). Segmentation and ΔF/F extraction were performed using the CaImAn pipeline, and only accepted components were included in subsequent analysis. For each segmented neuron, frequency, amplitude, and area under the curve were calculated using custom-developed code; the hyperactive neuron was defined as cells exhibiting 12 or more Ca^2+^ transients per minute. To assess network synchronization, the cross-correlation coefficient was computed for all neurons within each ROI.

For microglia surveillance analysis, each ROI data from *in vivo* 2P imaging was merged to a z-stack and motion corrected using Image Stabilizer and Poorman3D plugin. Maximum projection was generated from each time point, and then the cumulative maximum projections of the microglial image from time t and previous time points (t0, t0 + t1, t0 + t1 + t2, …) for every z-slice were generated. Microglia around the amyloid plaque were then cropped for further comparison. Microglia coverage for each image was calculated, and the cumulative volume filled by microglia every 2 min for 50 min was determined. The same threshold was used for all time points for each image. For microglia–neuron interaction analysis in awake mice, neuronal segmentation and ΔF/F signal extraction were performed using the CaImAn pipeline. Custom Python scripts were used to calculate the distance between each neuron and surrounding microglial pixels. The number of microglial pixels within 15 μm of each firing neuron was quantified. Analyses were restricted to neurons located within 50 μm of amyloid plaques.

For cell contact analysis, Z-stack images (20–40 μm) from single time-point in vivo two-photon imaging of anesthetized mice were analyzed using Imaris software (Bitplane). Imaging channels included GFP-labeled microglia, jRCaMP1b-expressing active neurons, and Methoxy-X04–labeled amyloid plaques. Image stacks were imported into Imaris, and 3D surface reconstruction was performed for each channel. Background subtraction and size filtering were applied to remove small disconnected particles (microglia, ~2–3 μm; neurons, ~3 μm; amyloid plaques, ~2 μm). To exclude dystrophic neurite signals associated with plaques, neuronal signals overlapping with amyloid plaque surfaces were removed from subsequent analyses. The cleaned surface objects were then used to quantify microglia–neuron contacts and to calculate intercellular distances between microglial processes and neuronal structures.

3DMorph was used to analyze microglial morphology in MATLAB. The GFP channel from the first time point of the surveillance period was extracted from *in vivo* 2P imaging data (512 × 512 pixels per image). Microglial clusters were identified using interactive mode for each image. Only microglial clusters located surrounding amyloid-β plaques (within 20 μm), were included in the analysis, with each cluster treated as a single object. Isolated microglia away from amyloid plaques were excluded from morphological analysis. 3DMorph automatically calculated territory volume, ramification index, number of endpoints, and total branch length for each microglial cluster.

Microglia motility was analyzed using ProMolJ. Image size reduction, pre-processing, selection of processes, process reconstruction, and motility analysis were performed using the default setting with the following parameter: Image corrections (Macro 3), M3.2, Max sigma=85. Process selection (Macro 4), M4.1, Threshold level = 6 or 7 depending on image quality. Motility analysis (Macro 5), M5.1, Time Interval = 0.5 min, M5.2 3D Gaussian sigma = 1, M5.3 Skeleton smoothing range = 2, M5.4, Maxima threshold = Same as M4.1. μm/pixel = 0.311701 Custom code for microglia spontaneous imaging was developed for this study and will be available on Github.

#### Single Nuclei RNA-seq

Mice were perfused with cold PBS and brains were then dissected and flash frozen. 50 mg tissue spanning cortex and hippocampus was homogenized in lysis Buffer (10 mM Tris-HCl, pH 7.4, 10 mM NaCl, 3 mM MgCl_2_, and 0.025% NP-40), and incubated on ice for 15 min. The suspension was filtered through a 70-μm filter to remove debris and pelleted at 300 xg for 10 min at 4 °C. Nuclei were washed and filtered twice with Nuclei Wash (1% BSA in PBS with 0.2 U μl^−1^ RNasin (Promega)). Using the Nuclei Isolation Kit (Sigma), nuclei were resuspended in 500 μl wash buffer and 900 µl 1.8 M sucrose. A gradient was prepared by layering 500 μl 1.8 M sucrose followed by centrifugation at 13,000 xg for 45 min at 4 °C. The nuclei pellet was resuspended in wash buffer at 1,000 nuclei μl^−1^ and filtered through a 20-μm filter.

For snRNA-seq analysis, a total of 14 samples were each loaded at target of 16,000 nuclei on a 10X Chromium Single Cell 3’ v3 kit (Rev C). Following manufactures instructions and QC parameters, libraries were then sequenced across three lanes of a Novaseq 6000. 10X Genomics cellranger suite was used to map and QC raw reads. The SoupX package (v1.6.2) was used for removal of ambient RNA contamination. Briefly, for each experimental sample, feature-barcode matrices from Cellranger were loaded with the function SoupX::load10X, and the degree of ambient RNA contamination was estimated using SoupX::autoEstCont, with default parameters; feature counts corresponding to ambient RNA were then removed using SoupX::adjustCounts, with the parameter roundToInt = T. The adjusted count matrices for all samples were then combined into a single count matrix and loaded into a Seurat object (Seurat v5.1.0) along with relevant experimental metadata for downstream pre-processing. After removal of ambient RNA contamination, initial downstream pre-processing consisted of computing Pearson residuals from regularized negative binomial regression as described in Hafemeister et al.,^99^ followed by dimensionality reduction and Leiden-based clustering according to the standard Seurat workflow. The following functions were sequentially applied to the initial Seurat object: 1) Seurat::SCTransform, with conserve.memory = T and do.correct.umi = F; 2) Seurat::RunPCA, with default parameters; 3) Seurat::FindNeighbors, with dims = 1:50; 4) Seurat::FindClusters, with resolution = 0.8; and 5) Seurat::RunUMAP, with dims = 1:50 and return.model = T. Due to the large number of cells in the dataset, cross-sample integration was not performed at this stage.

Following the initial pre-processing described above, the Azimuth package (v0.5.0)^100^ was used to generate automated cell type annotations based on alignment to a gold-standard reference dataset^101^, using the following function call: *Azimuth::RunAzimuth*, with *reference = ‘mousecortexref’*. In addition to predicted cell type labels (‘predicted.subclass’), Azimuth also generates a score reflecting the confidence for the label prediction (‘predicted.subclass.score’), as well as a score reflecting how well-represented the queried cell is in the reference dataset (‘mapping.score’). Since low quality cells or doublets are not present in the reference dataset, such cells in the query dataset tend to have low mapping scores, which was useful for downstream quality control.

For initial quality control, the cluster assignments, relevant experimental metadata, predicted cell type labels and mapping score from Azimuth, and QC metrics such as the total number of genes detected and the percent of mitochondrial transcripts were visualized with the UMAP of the entire dataset. Then clusters clearly corresponding to low quality cells (as judged by having lower than expected number of genes detected, a mixture of predicted cell type labels, and lower than expected mapping scores) or doublets (as judged by a lower than expected mapping scores and a mixture of predicted cell type labels) were removed. Following removal of such clusters, the pre-processing workflow described above was repeated, generating new cluster assignments, which were then manually assigned broad cell type annotations (glutamatergic neuron, GABAergic neuron, astrocyte, oligodendrocyte, oligodendrocyte precursor cell, microglia, endothelial cell, vascular-like meningeal cell, pericyte) based on the predominant Azimuth predictions (‘predicted subclass’) for each cluster.

Following initial quality control and manual broad cell type annotation, cells of each broad cell type were then analyzed separately in sub-clustering analysis. First, the same pre-processing workflow consisting of SCTransform followed by dimensionality reduction was applied, followed by visualization of subcluster assignments, relevant experimental metadata, QC metrics, and predicted subclass identities from Azimuth. Cells with predicted subclass identities not consistent with the broad cell type were removed, followed by removal of subclusters that corresponded to low-quality cells or doublets, using the same qualitative criteria as described above. Afterwards, cross-sample integration was performed for all broad cell types except for glutamatergic neurons and GABAergic neurons, which did not demonstrate any appreciable sub-clustering driven by sample identity and for which the same pre-processing workflow was applied after data cleaning. For all other broad cell types, when the minimum number of cells in the smallest cluster was greater than 50 in the sample with the fewest number of cells, cross-sample integration was achieved using *Seurat:: IntegrateLayers*, with *method = RPCAIntegration*, *normalization.method = ‘SCT’*, and *reference* set to one of the control samples; otherwise, cross-sample integration was achieved by setting *method = HarmonyIntegration.* Following cross-sample integration, subcluster assignments, relevant experimental metadata, QC metrics, and predicted subclass identities from Azimuth were visualized on the UMAP generated from the cross-sample integration, and residual doublet or low-quality clusters were removed. For broad cell types that consist of multiple subclasses (eg glutamatergic neurons), the sub-clusters were then manually assigned subclass identities based on the predominant Azimuth prediction for each sub-cluster (eg L2/3 IT).

Following quality control at the sub-clustering level, cross-condition differential expression analysis was performed on pseudobulked samples at the subclass level as follows: the data was converted to a *SingleCellExperiment* object *sce*, then pseudobulk samples were generated using *scater::aggregateAcrossCells* (scater v1.34.0), *with id = colData(sce)[, c(‘Sample_number”, “subclass.assignment”)]*, where the parameter *id* specifies the metadata columns according to which the pseudobulk aggregation should be performed. The pseudobulked SingleCellExperiment object was then filtered to remove pseudobulk samples containing fewer than 10 cells.

Differential expression analysis was then performed using the function *scran::pseudoBulkDGE* (scran v1.34.0), which is a wrapper function built around edgeR (v4.4.1).^102^ The design formula ~ Sex + Genotype was used, where Genotype is a factor (with the levels ‘WT’, ‘*Fgg^γ390–396A^’*, ‘5XFAD’, ‘5XFAD*:Fgg^γ390–396A^*’), and contrasts corresponding to the following comparisons were extracted: 1) ‘5XFAD’ vs ‘WT’, 2) ‘*Fgg^γ390–396A^*’ vs ‘WT’, and 3) ‘5XFAD: *Fgg^γ390–396A^*’ vs ‘5XFAD’. Due to the limited sensitivity of pseudobulked differential expression analysis with the sample sizes in our study,^103^ differentially expressed genes (DEGs) were called based on raw *P* values for downstream analyses.

#### Single-Nucleus and Single-Cell RNA-Sequencing Data Combination

Single-nucleus RNA-sequencing (snRNA-seq) (this paper) and single-cell RNA-sequencing (scRNA-seq) (Mendiola et al., ^45^) datasets were processed using the Seurat package (v5.1.0) ^104^ in R (v4.4.1). The raw counts of the snRNA-seq Seurat object (consisting of multiple cell types) were merged with the raw counts of the scRNA-seq dataset, which included only microglia. Normalization and variance stabilization were performed on the merged Seurat object using sctransform (vst.flavor = “v2”).^99,105^ Dimensionality reduction was conducted using principal component analysis (PCA), followed by Uniform Manifold Approximation and Projection (UMAP) using the top 10 principal components. The UMAP projection positioned microglia from scRNA-seq adjacent but not entirely overlapping with the microglia from snRNA-seq; however, due to the presence of only a single cell type in the scRNA-seq dataset, differences due to biological and technical (single cell vs single nuclei, different processing dates, library preparation operator) reasons between the two datasets remained difficult to ascertain and correct for.

To address dataset-specific variation, two integration methods were evaluated. SCTransform normalization while adjusting for (or regressing out) dataset-specific differences was applied, but this did not alter the UMAP projections. Additionally, Seurat’s Canonical Correlation Analysis (CCA) integration was tested but did not improve clustering by cell type. Therefore, the subsequent CellChat analysis was conducted using the merged, SCT-normalized dataset (without regressing out features).

#### Cell-Cell Communication Analysis Using CellChat

The CellChat (v2.1.2)^106^ analysis was conducted using the merged, SCT-normalized dataset (without regressing out features) to investigate intercellular communication differences between 5XFAD and 5XFAD: *Fgg^γ390–396A^* genotypes. Genotype-specific CellChat objects were created, and ligand-receptor interactions were inferred using CellChatDB.mouseV2 as the reference database. To characterize genotype-specific communication networks, overexpressed genes and interactions were identified for each genotype. Communication probabilities were computed using the triMean method, and networks were filtered to retain interactions present in at least 10 cells. Aggregated signaling networks were inferred, and circle plots were generated to visualize differences in the number and strength of interactions between genotypes.

To assess alterations in intercellular communication, the compareInteractions function was used to quantify differences in the number and strength of ligand-receptor interactions between 5XFAD and 5XFAD:*Fgg^γ390-396A^*. Network visualization was conducted using the netVisual_diffInteraction function, which generates circle plots highlighting interactions that were increased or decreased between genotypes. To assess cell-type-specific communication patterns, the netVisual_heatmap function was used to visualize differences in the number and strength of interactions in the inferred cell-cell communication network between the two genotypes. Heatmaps were generated to compare interaction patterns across cell types, and bar plots were used to summarize incoming and outgoing signaling activity. Additionally, the overall information flow within signaling pathways between genotypes was evaluated, with pathways ranked based on differences in inferred network activity and genotype-specific enrichment visualized using bar plots. CellChat objects from different genotypes were then merged to compare interaction changes across genotypes. To visualize microglia-neuron interaction changes, ligand-receptor pairs were analyzed, with ligands from microglia and receptors from neurons (both excitatory and inhibitory) used for generating bubble plots and Chord diagrams.

#### Correlative light-electron microscopy (CLEM)

After completing *in vivo* 2P imaging, mice were anesthetized via intraperitoneal injection of ketamine/xylazine and transcardially perfused with a brief flush of Ringer’s solution containing heparin and xylocaine, followed by approximately 80 ml of 0.5% glutaraldehyde and 4% paraformaldehyde in sodium cacodylate buffer. The brains were post-fixed overnight and sectioned at 150 μm thickness. The slices were then stained for 1 hour on ice with DRAQ5 (Biostatus) diluted 1:1,000 in cacodylate buffer, followed by three washes in cacodylate buffer. The areas of interest were identified and confocal microscopy volumes were acquired using an Olympus FluoView 1000 inverted microscope (20× and 60× water objectives).

The slices were post-fixed overnight at 4 °C in cacodylate buffer containing 2.5% glutaraldehyde, followed by washing in cacodylate buffer containing 100 mM glycine to quench excess aldehydes. The slices were stained with 2% reduced osmium tetroxide, 0.05% thiocarbohydrazide, 2% osmium tetroxide, 2% uranyl acetate, and Walton’s lead aspartate solution, with thorough double-distilled water washes between each step. The sections were subsequently dehydrated through a graded ethanol series (70%, 90%, 100%) and dry acetone solutions, followed by infiltration and embedding in Durcupan ACM resin. The resin-embedded samples were then cured at 60°C for 48 hours.

The slices were mounted on the end of a small aluminum pin, and low-resolution XRM volumes were collected at 80 kV using a Zeiss Versa 510 XRM system. The vasculature pattern was used as a landmark for localizing the region of interest (ROI) within the embedded section. The blocks were then trimmed to < 1 mm × 1 mm and mounted onto SBEM specimen rivets using conductive silver epoxy (Ted Pella), followed by overnight curing at 60°C. After further trimming using an ultramicrotome, higher-resolution XRM scans were performed to precisely target the SBEM imaging stage to the ROIs.

For SBEM data acquisition, 2 × 2 montage volumes were collected using a Zeiss Gemini 300 SEM equipped with a Gatan 3View and OnPoint backscatter detector system. The SBEM volumes were acquired at 2.5 kV EHT with a pixel resolution of 6.5 nm in XY, a Z-step size of 60 nm, and a dwell time of 1 µs per pixel. To generate the final SBEM volumes, the ‘tiltxcorr’ program in IMOD^94^ was used to apply cross-correlation alignment, eliminating minor inter-slice jitter. SBEM and confocal datasets were co-registered using the ‘Landmark Image Warp’ module in Amira (2020.2 version; ThermoFisher Scientific), aligning nuclear features and

#### SBEM Semi-automatic Segmentation

For semi-automated 3D reconstruction of microglia-neuron contacts, we applied CDeep3M,^107^ a deep-learning-based image segmentation tool for large-scale microscopy datasets. A pre-trained model (https://doi.org/10.7295/W9CDEEP3M50692) optimized for plasma membrane detection in SBEM image volumes was used to segment the entire SBEM dataset. This model had been previously trained on ground-truth annotations from other SBEM image volumes. The CDeep3M-generated cell boundary predictions and raw SBEM image volumes were subsequently imported into Volume Annotation and Segmentation Tool (VAST) Lite software,^95^ where neurons and microglia were rapidly traced using machine-assisted boundary detection as a guide. This AI-assisted annotation enabled high-throughput 3D reconstruction of microglia-neuron interactions at the nanoscale, facilitating precise quantification of microglial process morphology and contact frequency with neuronal compartments.

#### CSF protein analyses Somascan

The dataset used for the selection of CSF samples was previously described extensively^108^. Study protocols were approved by their respective Institutional Review Boards. Research was performed in accordance with the Code of Ethics of the World Medical Association. Written informed consent was obtained from all patients before data collection. Proteomic data were generated from CSF using the SomaLogic Proteomics Platform. The raw data underwent quality control, calibration, and normalization steps. From the proteomics data, proteins that passed QC were selected to test the association between fibrinogen levels in linear models. Proteins that were significantly correlated with fibrinogen (adjusted P value < 0.05) were used for pathway analysis with ClusterProfiler. For cell type enrichment analysis, the expression profile of the healthy brain was obtained from the Human M1 10x database on the Allen Institute. Each gene correlated with fibrinogen was given 1 point and distributed to each cell type based on their gene expression, and the total enrichment score for each cell type is the sum of points from all fibrinogen-correlated genes. Genes not expressed in the brain based on the human M1 10x database were not included in the analysis.^108^

### QUANTIFICATION AND STATISTICAL ANALYSIS

All values are reported as mean ± s.e.m. Unpaired t-tests were used to compare between two groups. For comparisons involving more than two groups, one- or two-way ANOVA followed by Tukey’s post hoc test for multiple comparisons was used. For analysis of multiple timepoint and multiple genotypes, generalized linear mixed model was used with Bonferroni post-hoc tests to account for multiple comparisons. For all *in vivo* experiments, mice were randomized and experiments were conducted in a blinded manner to the mouse genotype and antibody administration. Genotype and treatment assignment were revealed after image quantification.

---

**Figure S1.**
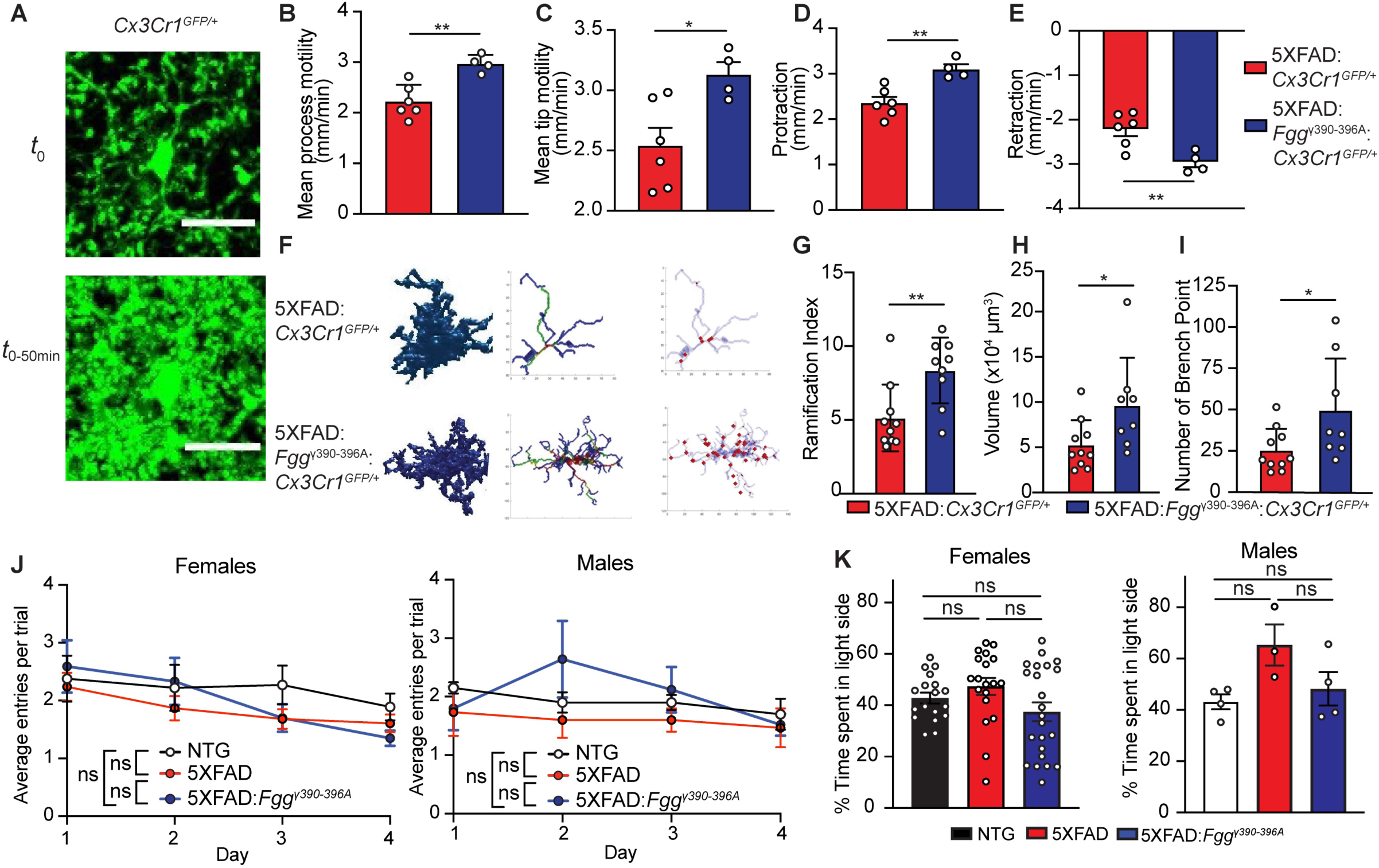
Fibrin abrogates microglia motility in 5XFAD mice. (**A**) Representative of *in vivo* 2P time-lapse imaging of microglia surveillance in *Cx3CR1^GFP/+^* mice. (**B-E**) Quantification of microglia motility surrounding amyloid plaque. Mean process motility(B), mean tip motility (C), protraction (D) and retraction (E). Data represents mean ± s.e.m from *n* = 6 5XFAD, n = 4 5XFAD: *Fgg^γ390-396A^*. *\*P* <0.05, by student t-test. (**F-I**) Analysis of the morphology of amyloid plaque in 5XFAD and 5XFAD: *Fgg^γ390-396A^* mice by 3DMorph. Representative image (F) and quantification of ramification index (G), skeleton (H) and branch point (I) of microglia cluster surrounding amyloid plaque analyzed by 3DMorph. Data represents mean ± s.e.m from *n* = 10 5XFAD, n=8 5XFAD: *Fgg^γ390-396A^*. *\*P* <0.05, **P <0.01, by student t-test. (**J**) Quantification of average entry per trial over 4 days in the Cost-Benefit Conflict (CBC) test. Data represents mean ± s.e.m from n = 13 NTG, n = 21 5XFAD, and n = 17 5XFAD:*Fgg^γ390-396A^* female mice and n = 4 NTG, n = 3 5XFAD, and n = 5 5XFAD:*Fgg^γ390-396A^* male mice, ns. ns, not significant. (**K**) Light/Dark preference determined by the percentage of time spent in the light side in the CBC test. Data represents mean ± s.e.m from n = 13 NTG, n = 21 5XFAD, and n = 17 5XFAD:*Fgg^γ390-396A^* female mice and n = 4 NTG, n = 3 5XFAD, and n = 5 5XFAD:*Fgg^γ390-396A^* male mice, ns. ns, not significant.

**Figure S2.**
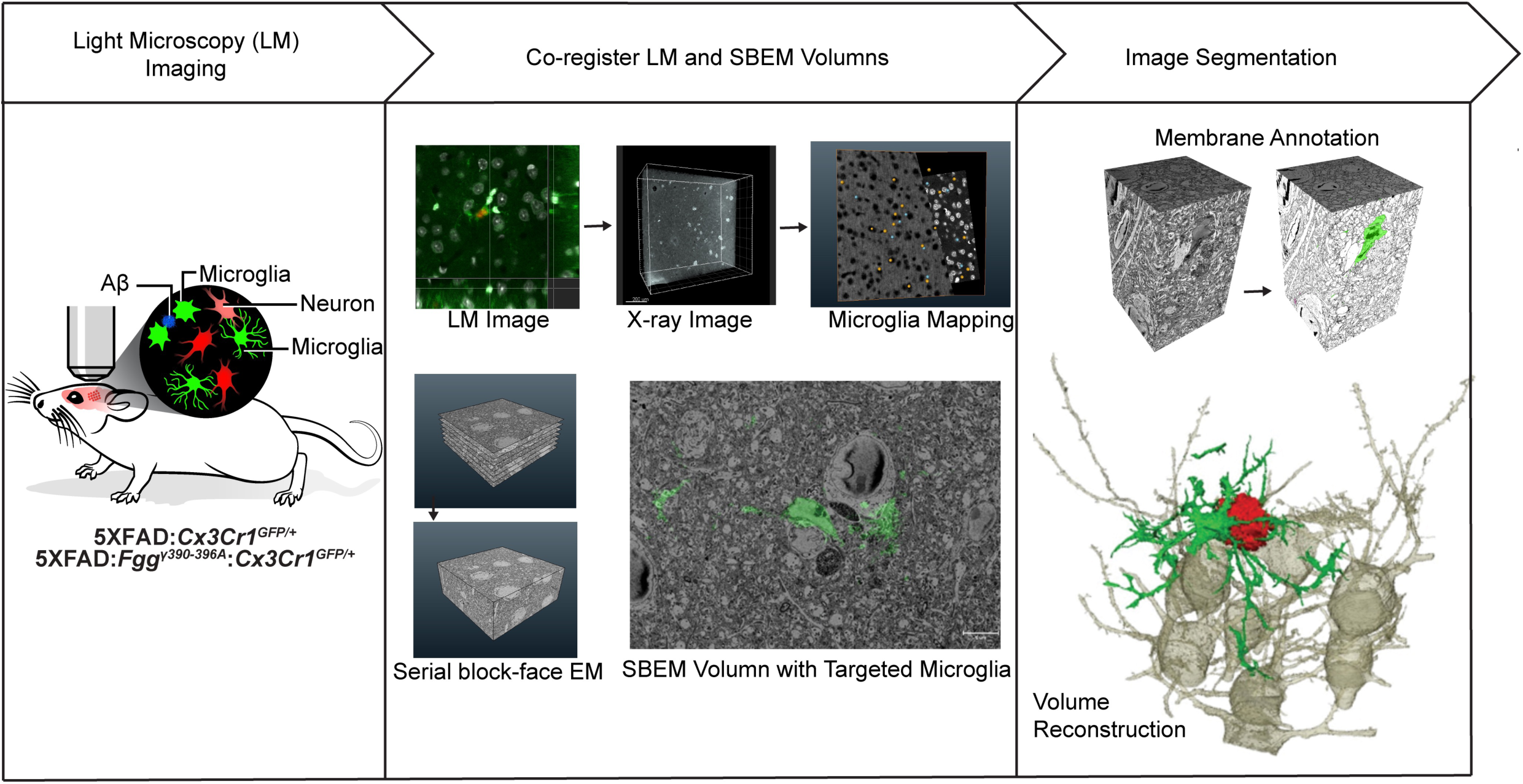
Multiscale multimodal imaging pipeline for microglia-neuron interaction. Schematic of multiscale, multimodal imaging pipeline to co-register LM and SBEM volumes for volumetric image analysis of microglial cell contacts with amyloid plaque and active neurons at sites of vascular disruption in 5XFAD mice.

**Figure S3.**
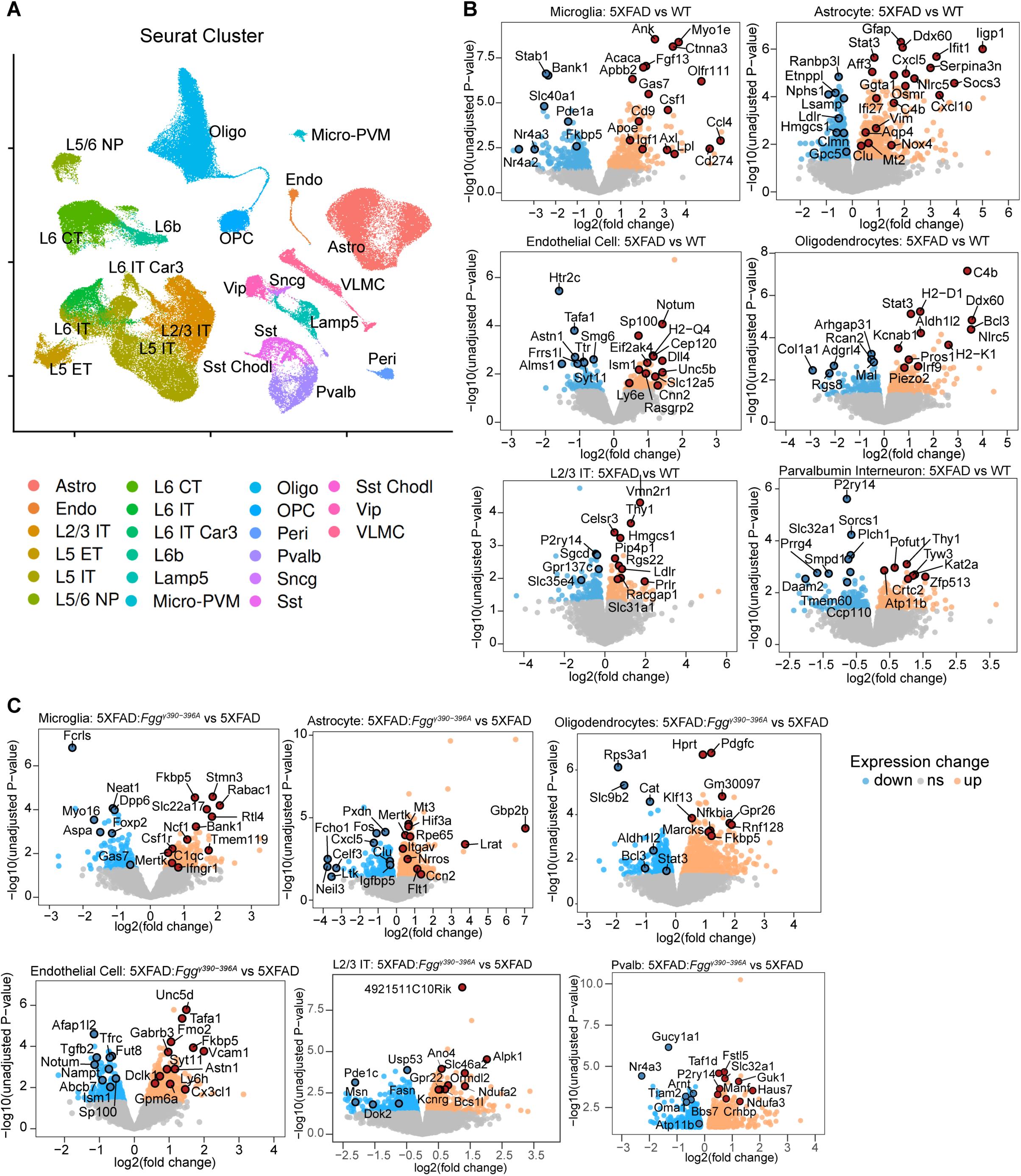
Differentially regulated genes identified by snRNA-seq in 5XFAD, 5XFAD*:Fgg^γ390-396A^* and littermate controls. (**A**) UMAP plot of 165,252 nuclei showing 21distinct cell types from 6-month old brains of WT, *Fgg^γ390-396A^*, 5XFAD and 5XFAD:*Fgg^γ390–396A^* mice. Nuclei are colored by the identified cell type clusters and displays aggregated samples from *n* = 3 WT, n = 3 *Fgg^γ390-396A^*, n = 4 5XFAD and *n* = 4 5XFAD:*Fgg^γ390–396A^* mice. (**B**) Volcano plots of DEGs in each cluster between 5XFAD versus WT mice of each cell type by pseudobulk analysis. Dots depict average log_2_ fold change and −log10 unadjusted P values for each gene colored by significance cutoff (log_2FC_ > 1.5, unadjusted P value < 0.05). Data represent *n* = 3 WT, n = 3 *Fgg^γ390-396A^*, n = 4 5XFAD and *n* = 4 5XFAD:*Fgg^γ390–396A^* mice. (**C**) Volcano plots of DEGs in each cluster between 5XFAD*:Fgg^γ390-396A^* versus 5XFAD mice of each cell type by pseudobulk analysis. Dots depict average log_2_ fold change and −log10 unadjusted P values for each gene colored by significance cutoff (log_2FC_ > 1.5, unadjusted P value < 0.05). Data represent *n* = 3 WT, n = 3 *Fgg^γ390-396A^*, n = 4 5XFAD and *n* = 4 5XFAD:*Fgg^γ390–396A^* mice.

**Figure S4.**
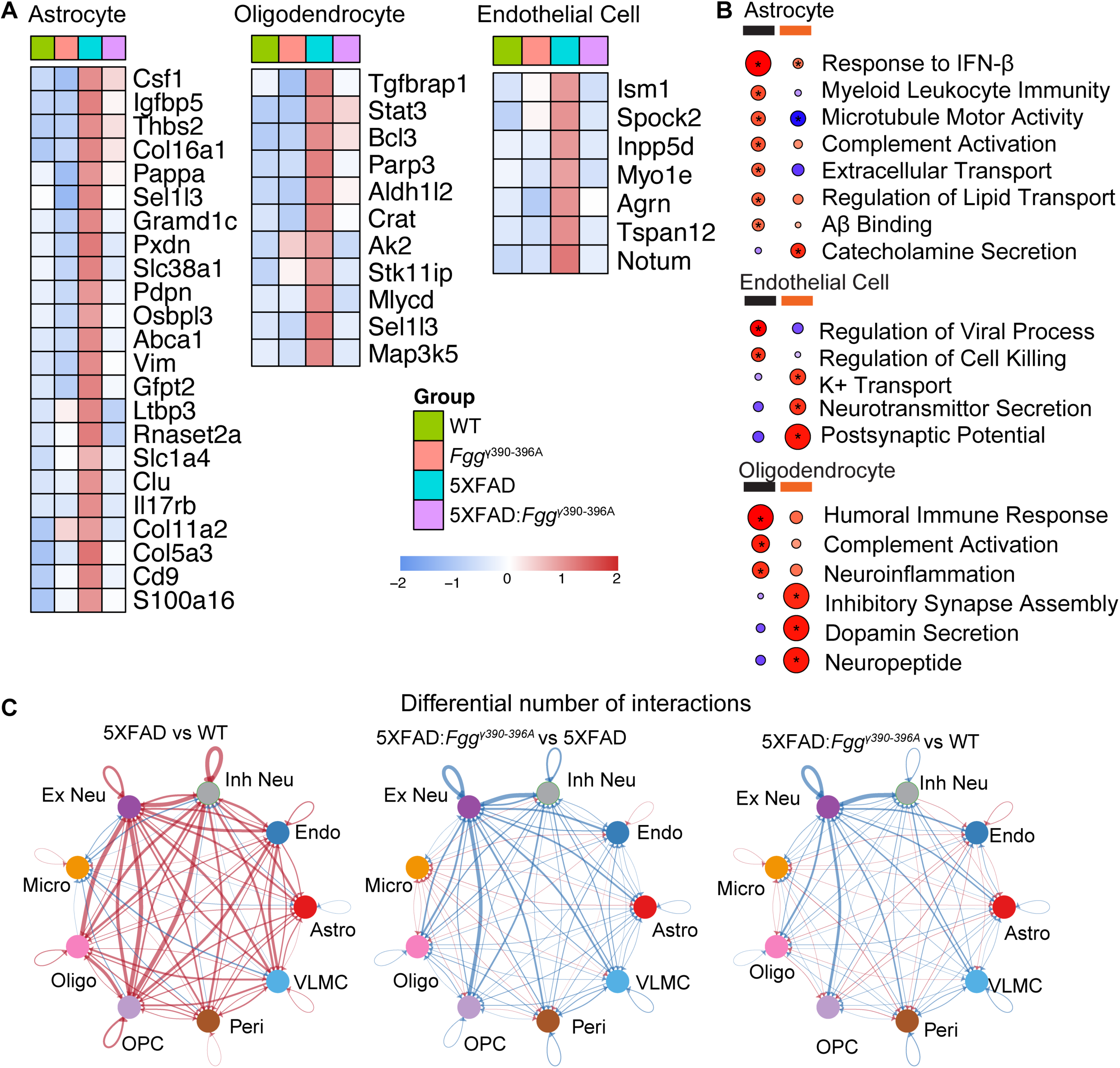
Fibrin-regulated glial and endothelial cell gene signatures in AD mice. (**A**) Heatmap of significant DEGs in the Astrocytes, oligodendrocytes, and endothelial cells from WT, *Fgg^γ390-396A^*, 5XFAD, and 5XFAD:*Fgg^γ390–396A^* mice by snRNA-seq (*P* < 0.05 in Pseudobulk analysis by 5XFAD vs WT and 5XFAD:*Fgg^γ390-396A^* vs WT). Colors indicate the average scaled expression level for each genotype. Data points are averages of *n* = 3 WT, n = 3 *Fgg^γ390-396A^*, n = 4 5XFAD and *n* = 4 5XFAD:*Fgg^γ390–396A^* mice. (**B**) Gene Set Enrichment Analysis (GSEA) of snRNA-seq data comparing 5XFAD vs WT and 5XFAD:*Fgg^γ390–396A^* vs 5XFAD in astrocyte, endothelial cell and oligodendrocyte clusters using gene list ranked by log2FC from Pseudobulk analysis. (**C**) Circle plots showing total strength of cell-cell interactions differentially regulated between genotype comparisons by Cellchat analysis. The edge size depicts the strength of interactions. The nodes depict cell clusters as identified in the snRNA-seq dataset shown in Figure 3.

**Figure S5.**
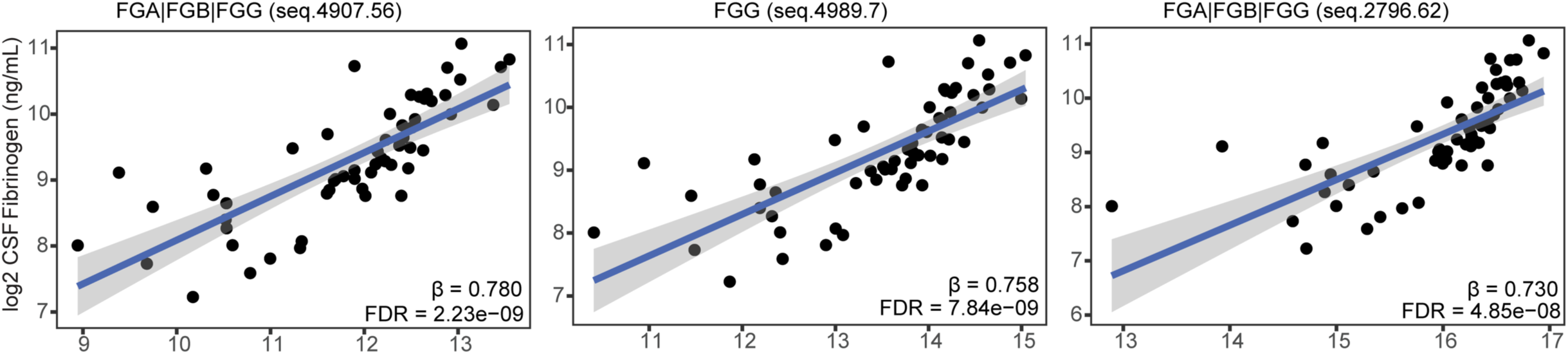
CSF fibrinogen levels measured by ELISA correlate with SomaScan measurements. Dot plots show the correlation between CSF fibrinogen levels determined by ELISA and fibrinogen levels measured by three SomaScan aptamers (seq.4907.56, seq.4989.7, and seq.2796.62).

**Supplementary Video 1** Time-lapse in vivo 2P imaging of cumulative volume sampling of microglia surveillance in *Cx3cr1*^GFP/+^, 5XFAD:*Cx3cr1*^GFP/+^ and 5XFAD: *Fgg^γ390-396A^*:*Cx3cr1*^GFP/+^ mice. A 50-µm *z*-stack was acquired in the cortex every 2 min for 50 min. Each frame is an overlay of the maximum-intensity projection acquired at *t_i_* and all previous frames. Scale bar, 20 µm.

**Supplementary Video 2** 3D reconstruction of cortical microglia, neuron and Aβ plaque by SBEM and CDEEP3M machine-deep-earning volumetric image analysis in 5XFAD:*Cx3cr1*^GFP/+^ mice. A representative GFP+ microglia (green) around an amyloid plaque (red), neighboring neurons (yellow) and distal neurons (grey).

**Supplementary Video 3** 3D reconstruction of cortical microglia, neuron and Aβ plaque by SBEM and CDEEP3M machine-deep-earning volumetric image analysis in 5XFAD: *Fgg^γ390-396A^*:*Cx3cr1*^GFP/+^ mice. A representative GFP^+^ microglia (green) around an amyloid plaque (red), neighboring neurons (yellow) and distal neurons (grey).

**Table S1.** DEG from snRNA-seq for each celltype by pseudobulk analysis, related to Figure 3

**Table S2.** Significantly changed pathways of each cell type from the snRNA-seq dataset by gene set enrichment analysis, related to Figure 3

**Table S3.** CellChat-predicted microglia–neuron ligand–receptor interactions across genotypes, related to Figure 3

**Table S4.** CSF proteomic dataset from Alzheimer’s disease patients used for correlation analysis with CSF fibrinogen level measured by ELISA, related to Figure 4

**Table S5.** CSF SOMAmers significantly correlated with CSF fibrinogen levels measured by ELISA, related to Figure 4

